# Comparisons among Yanomami populations reveal transitional gut microbiome configurations under urban influence

**DOI:** 10.64898/2026.10.27.762328

**Authors:** Lucas Amoroso Lopes de Carvalho, Marie Thorp, Maarten Ciers, Chris Callewaert, David Good, Emma Allen-Vercoe, Lars Vereecke, Luciana Campos Paulino

## Abstract

Urbanization influences gut microbiome composition and function, yet comparisons between contrasting lifestyles offer limited insight into which microbial features are most sensitive to increasing urbanization. We analyzed gut shotgun metagenomes from Yanomami participants, an Indigenous Amazonian people who maintain traditional livelihoods, across three geographically distinct populations with varying degrees of urban influence, as well as an urban Belgian cohort. Belgian and Yanomami microbiomes differed markedly in taxonomic composition and functional potential. The comparison between Bicho-Açu, the most urban-influenced Yanomami population, and Xamakorona, the least urban-exposed, revealed that although Bicho-Açu presented a broadly Yanomami-associated microbiome, it exhibited selected urban-associated features, including greater *Bacteroides* abundance and a higher relative abundance of genes involved in glycosaminoglycan degradation and secondary bile acid biosynthesis pathways. Furthermore, genus-level co-occurrence network connectivity showed an orderly shift across populations, declining progressively with increasing urban influence. Carbohydrate-active enzyme profiles further showed stronger non-starch-polysaccharide-associated signals in Belgium and resistant-starch-associated features in Xamakorona, suggesting distinct carbohydrate-processing strategies. Our findings support a model in which urbanization reshapes microbial composition, their ecological interactions, and metabolic capacity, with transitional configurations emerging through selective shifts across these dimensions.

## Introduction

Urbanization is rapidly transforming the ecological conditions under which human gut microbial communities are acquired and maintained. Changes in diet and food processing, antibiotic use, sanitation, personal hygiene, and contact with environmental microorganisms might simultaneously alter microbial acquisition through vertical or horizontal transmission, as well as the microbial substrates and ecological niches available within the intestine^1–3^. These changes are biologically relevant because gut microorganisms transform dietary and host-derived molecules into bioactive compounds that influence intestinal, immune, and metabolic physiology^4,5^. Therefore, fully understanding microbiome urbanization requires determining not only which organisms are supported or depleted but also how changing exposures reorganize microbial communities and their encoded metabolic functions.

Comparisons between urbanized and non-urbanized populations have established noticeable differences in gut microbiome composition and functional potential across diverse geographic settings^6–9^. Urbanization is frequently associated with reduced richness and prevalence of particular microbial lineages, including specialized plant-polysaccharide degraders, together with the expansion of taxa capable of exploiting animal-derived nutrients and readily accessible dietary or host glycans^10–13^. However, urbanization-associated remodeling cannot be reduced to generalized diversity loss. It also encompasses abundance shifts among taxa that remain widely shared and the reorganization of metabolic repertoires^3,9,14^. These endpoint differences are now well documented; what remains less clear is how they emerge as populations experience differing degrees and forms of urban influence.

Most comparative studies contrast populations that already occupy widely separated lifestyle, geographic, genetic, and socioeconomic contexts. Such study designs do not readily allow determination of whether the various dimensions of the microbiota, such as taxonomic and functional composition, as well as the structure of associations between taxa, change together or respond differently. Intermediate states of urbanization could therefore represent coherent community-wide shifts, mosaics combining microbial features associated with hosts experiencing different degrees of urban influence, or stable adaptations specific to particular populations. Studies along rural-urban and lifestyle gradients increasingly indicate that microbiome responses are heterogeneous and contingent on local ecology, geography, and population history^3,14–17^. Comparisons among related populations experiencing different urban-associated exposures can reduce some of the heterogeneity inherent to distant and broad lifestyle contrasts and help establish which microbiome features are most sensitive to urbanization-associated change, which respond only to greater exposure levels, and which remain comparatively stable.

The Yanomami provide a rare opportunity to address this question. They are an Indigenous people inhabiting the Brazilian and Venezuelan Amazon. Many maintain traditional livelihoods based on communal living, with practices that include subsistence horticulture, hunting, fishing, and gathering^18^. Their communities differ substantially in geographic accessibility and contact with non-Indigenous society, resulting in uneven exposure to urban centers, industrialized foods, personal hygiene products, healthcare, and antibiotics. Previous studies have documented distinctive and highly diverse Yanomami gut microbiomes and demonstrated variation among villages^11,19,20^. In this context, three related Yanomami subpopulations experiencing different degrees of urban influence provide an informative internal comparison alongside a strongly contrasting urban cohort. Additionally, profiling a substantial Yanomami cohort is important because Indigenous populations remain markedly underrepresented in global metagenomic resources. This underrepresentation limits current understanding of the diversity and population-specific characteristics of human gut microbial ecology^18,21,22^.

Here, we used shotgun metagenomics to characterize the gut microbiomes of 78 individuals from three Yanomami subpopulations along an urbanization gradient and compared them with 39 participants from an urban Belgian cohort. We integrated taxonomic composition with genus-level co-occurrence networks and functional profiling of KEGG orthologs and metabolic pathways. We also examined genes associated with microbial butyrate biosynthesis pathways and characterized carbohydrate-active enzyme repertoires, including their predicted substrate assignments. This approach allowed us to distinguish broad Belgian-Yanomami differences from changes within the Yanomami populations and identify features associated with increasing urban influence. By connecting these patterns to microbial substrate use and metabolic capacities, our study also provides a basis for testing dietary and microbiota-directed strategies that target specific microbial guilds and functions to support host health.

## Results

We analyzed 117 fecal shotgun metagenomes in a cross-sectional comparison of an urban Belgian cohort (BEL; n = 39) and three Indigenous Yanomami subpopulations living along the Marauiá River in Amazonas, Brazil: Bicho-Açu (BIC; n = 31), Curuá (CUR; n = 20), and Xamakorona (XAM; n = 27) (**Fig. 1a**). The Yanomami subpopulations differ in geographic remoteness and urban influence, according to a score that incorporates access to industrialized foods, personal hygiene products, and healthcare services, as well as previous antibiotic exposure^23^. BIC is closest to urban centers and has the greatest urban influence, CUR occupies an intermediate position, and XAM is the most remote and least urban-exposed. Participants included both sexes and ranged from childhood to older adulthood; demographic characteristics, urbanization scores, and geographic reference coordinates are provided in **Supplementary Table 1**.

**Figure 1.**
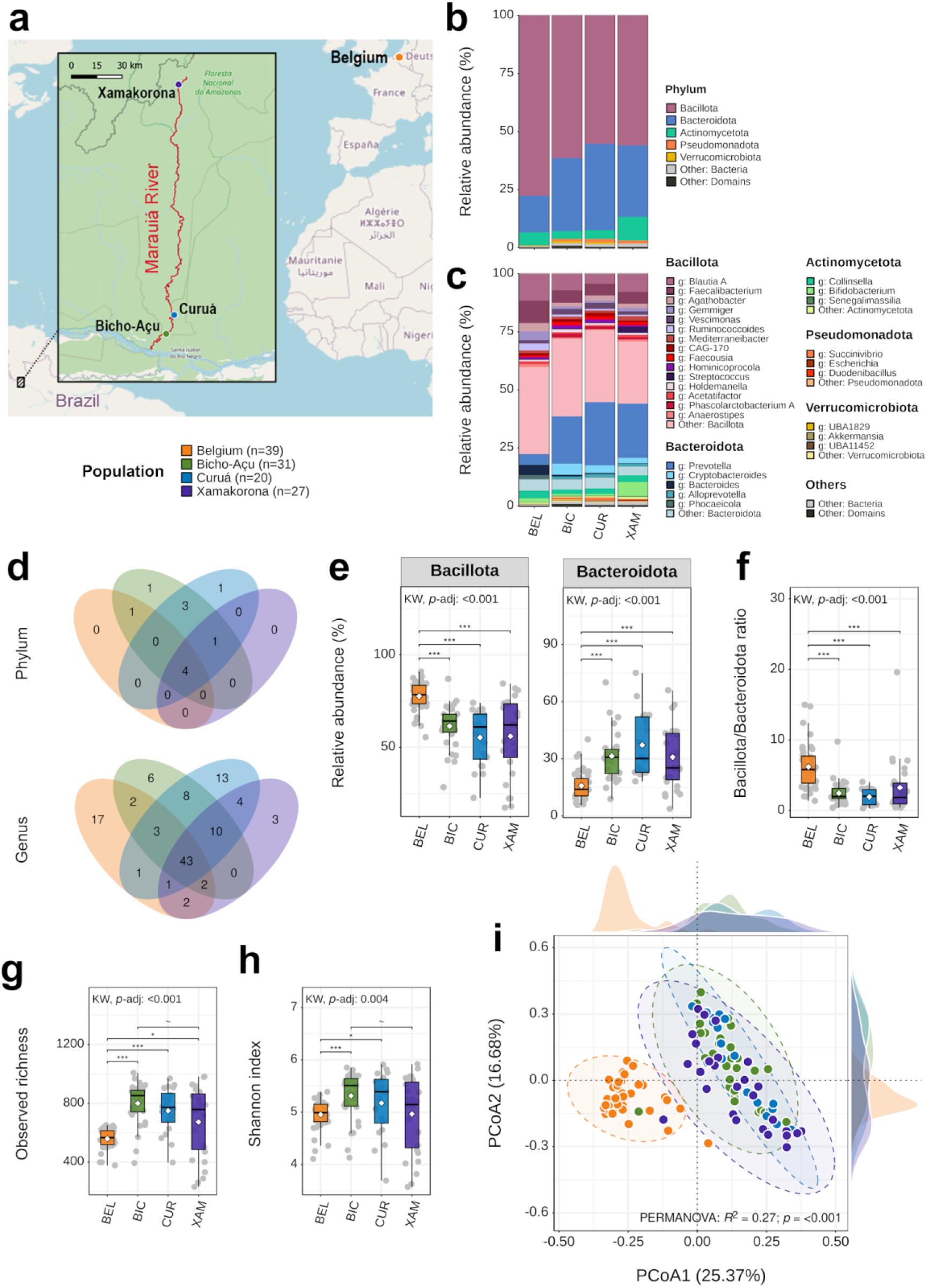
Taxonomic composition and diversity primarily distinguish Belgian and Yanomami gut microbiomes across the urbanization gradient. **a,** Geographic reference locations of the study populations in Belgium and the Brazilian Amazon. The inset shows the Yanomami subpopulations of Bicho-Açu, Curuá and Xamakorona along the Marauiá River in Amazonas state, Brazil. **b,** Mean relative abundance at the phylum level. The five most abundant phyla are shown individually; lower-abundance bacterial phyla and non-bacterial taxa are grouped as “Other: Bacteria” and “Other: Domains”, respectively. **c,** Mean relative abundance at the genus level, with genera color-coded by phylum. To preserve representation across dominant phyla, the 15 most abundant Bacillota genera, five most abundant Bacteroidota genera, and three most abundant genera from Actinomycetota, Pseudomonadota, and Verrucomicrobiota are shown. Remaining or genus-unclassified taxa are aggregated within their respective phyla or as “Other: Bacteria” or “Other: Domains”. **d,** Venn diagrams showing phyla and genera shared among or exclusive to populations; taxa were considered present when detected in ≥50% of individuals within a population. **e,** Relative abundance of Bacillota and Bacteroidota. Mean ± SD values were, respectively, BEL: 77.61 ± 7.87% and 15.83 ± 7.62%; BIC: 61.42 ± 11.67% and 31.45 ± 12.50%; CUR: 55.26 ± 14.76% and 37.26 ± 16.69%; XAM: 55.95 ± 20.91% and 30.87 ± 16.39%. Overall differences were significant for Bacillota (Kruskal–Wallis *H* = 46.90, FDR-adjusted *P* < 0.001) and Bacteroidota (H = 41.26, FDR-adjusted P < 0.001). **f,** Bacillota/Bacteroidota relative-abundance ratio. Mean ± SD: BEL, 6.20 ± 3.24; BIC, 2.47 ± 1.71; CUR, 1.94 ± 1.13; XAM, 3.26 ± 3.93 (Kruskal–Wallis *H* = 43.60, FDR-adjusted *P* < 0.001). **g,** Observed taxonomic richness: BEL, 558.33 ± 69.70; BIC, 799.52 ± 141.30; CUR, 749.50 ± 152.84; XAM, 672.67 ± 234.98 (mean ± SD; *H* = 38.01, FDR-adjusted *P* < 0.001). **h,** Shannon taxonomic diversity: BEL, 4.95 ± 0.29; BIC, 5.31 ± 0.46; CUR, 5.17 ± 0.61; XAM, 4.97 ± 0.72 (mean ± SD; *H* = 14.30, FDR-adjusted *P* = 0.004). For **e–h,** significant omnibus tests were followed by pairwise two-sided Wilcoxon rank-sum tests with Benjamini–Hochberg FDR correction. **i,** Principal coordinates analysis (PCoA) of Bray–Curtis dissimilarities calculated from taxonomic profiles. Density plots show the distribution of sample scores along PCoA1 (top) and PCoA2 (right), grouped according to the color coding in each panel. Population explained 27% of compositional variation (PERMANOVA: R2 = 0.27, F = 13.62, *P* < 0.001; 999 permutations). Pairwise PERMANOVA results are reported in **Supplementary Table 4**. Significance for FDR-adjusted pairwise comparisons is indicated as \*\*\**P* < 0.001, \*\**P* < 0.01, \**P* < 0.05, and ∼*P* < 0.1.

Metagenomic data recovery was consistently high: 90.16–92.52% of reads mapped to the corresponding population co-assembly at mapping quality ≥Q20 (**Supplementary Table 2**). Taxonomic classification rates also occupied narrow ranges across populations, spanning 96.72–97.17% at phylum level, and 81.28–85.09% at genus level (**Supplementary Fig. 1a**), providing broadly comparable taxonomic resolution for subsequent population comparisons.

### Belgian and Yanomami microbiomes differ in taxonomic composition

Bacteria dominated every population, accounting for 99.09–99.72% of sequences before abundance and prevalence filtering (**Supplementary Fig. 1b**). Archaea, Eukaryota and viruses each remained below 1% in every group, although their relative abundances differed among populations (Kruskal–Wallis *H* = 10.71–37.02, all FDR-adjusted *P* ≤ 0.013). At phylum level, the principal contrast involved Bacillota and Bacteroidota (**Fig. 1b,e**). Bacillota represented 77.61% of the BEL microbiome, compared with 61.42% in BIC, 55.26% in CUR and 55.95% in XAM. Bacteroidota showed a reciprocal distribution, increasing from 15.83% in BEL to 31.45% in BIC, 37.26% in CUR, and 30.87% in XAM, resulting in considerable differences in the ratio of these phyla between Belgian and Yanomami populations (**Fig. 1f**). Actinomycetota were particularly abundant in XAM (10.12%, versus 3.38–5.61% in the other populations), reflecting high relative abundances of *Bifidobacterium* in a subset of XAM samples (Supplementary Fig. 1c,d; population mean, 6.02%, versus 0.64–1.79% in the other populations).

The phylum-level differences were accompanied by changes across several abundant genera (**Fig. 1c**). Among Bacillota, *Blautia A* averaged 11.60% in BEL and 4.22–7.65% in the Yanomami subpopulations; *Faecalibacterium*, 9.62% and 5.15–5.36%; *Gemmiger*, 3.80% and 0.85–1.47%; and *Ruminococcoides*, 2.97% and 0.57–0.81%, respectively. Within Bacteroidota, *Prevotella* increased from 4.55% in BEL to 20.37–27.24% across BIC, CUR and XAM, whereas *Bacteroides* decreased from 4.39% to 0.25–0.78%. *Cryptobacteroides* showed a further Yanomami-associated pattern, averaging only 0.14% in BEL but 2.34–4.73% in the three Yanomami subpopulations.

Prevalence-based profile analysis nevertheless revealed a shared taxonomic core (**Fig. 1d**). Four phyla, namely Actinomycetota, Bacillota, Bacteroidota, and Pseudomonadota, occurred in at least half of the individuals in every population. Cyanobacteriota met this threshold in all three Yanomami subpopulations but not in BEL, whereas Bigyra and Spirochaetota were that prevalent only in BIC and CUR, respectively. At genus level, 43 taxa were prevalent across all four populations, including *Prevotella*, *Bacteroides*, *Blautia A*, *Faecalibacterium*, *Agathobacter*, *Roseburia* and *Coprococcus*. A further ten genera, including *Alloprevotella*, *Catenibacterium*, *Cryptobacteroides*, *Phascolarctobacterium A* and *Lentihominibacter*, were prevalent in each Yanomami subpopulation but not in BEL. Seventeen genera met the prevalence threshold only in BEL, six only in BIC, 13 only in CUR and three only in XAM. These designations indicate a core microbiota that varies across populations, showing a rearrangement between the urban population and the Yanomami, as well as internal variations within their subpopulations.

Alpha-diversity measures distinguished the populations in different ways (**Fig. 1g,h**). Observed richness was lowest in BEL and highest in BIC, with intermediate values in CUR and in XAM. Shannon diversity was highest in BIC, followed by CUR, whereas BEL and XAM had similar means (*H* = 14.30, FDR-adjusted *P* = 0.004). XAM also showed the widest within-population spread in Bacillota abundance, richness and Shannon diversity, consistent with the visibly heterogeneous individual profiles in **Supplementary Fig. 1c,d**. Despite these different alpha-diversity configurations, overall taxonomic composition was strongly structured by population (PERMANOVA: R^2^ = 0.27, *F* = 13.62, *P* < 0.001; **Fig. 1i**). All pairwise contrasts were significant, with the largest separations involving BEL (pairwise *F* = 22.14–24.44, *P* < 0.001) and smaller effects among Yanomami subpopulations (pairwise *F* = 2.62–4.66, *P* = 0.012 to <0.001; **Supplementary Table 4**). This population structure was not detectably modified by sex or age group, as neither the population-by-sex interaction (R^2^ = 0.018, *F* = 0.943, *P* = 0.519) nor the population-by-age-group interaction (R^2^ = 0.062, *F* = 0.869, *P* = 0.840) was significant (**Supplementary Fig. 1e,f**).

### Genus-level markers emphasize the Belgium–Yanomami contrast while distinguishing Yanomami subpopulations

MaAsLin2 identified 48, 64, and 45 differentially abundant genera in the BEL–BIC, BEL–CUR and BEL–XAM comparisons, respectively (**Fig. 2a**). Several highly abundant Bacteroidota taxa showed divergent associations in the Belgian–Yanomami contrast. *Prevotella* was the most enriched taxon overall in Yanomami relative to BEL, along with *Cryptobacteroides* and *Alloprevotella*. Conversely, *Bacteroides* was enriched in BEL, as were *Phocaeicola* and *Alistipes*. Prevalence further helps determine whether associations reflect changes in abundance or population occupancy. *Prevotella* was detected in every individual in all four populations, and *Bacteroides* was also nearly universal (100% in BEL, BIC, and CUR; 92.6% in XAM); their differential associations therefore primarily reflected abundance. By contrast, *Succinivibrio*, *Sodaliphilus*, and *Oribacterium* were not detected in BEL, occurred in only 19.4–29.0% of BIC participants, and exceeded 60% prevalence in both CUR and XAM. *Alistipes* showed the opposite occupancy pattern, decreasing from 94.9% prevalence in BEL to 32.3% in BIC, 15.0% in CUR, and was undetected in XAM. The taxonomic separation, therefore, combined the rebalancing of the abundances of certain widely shared genera with differences in population occupancy. The BIC subpopulation exhibited an intermediate occupancy pattern regarding discriminating genera, combining a lower prevalence of taxa associated with the Yanomami with a higher prevalence of taxa associated with the BEL, compared to the CUR and XAM groups.

**Figure 2.**
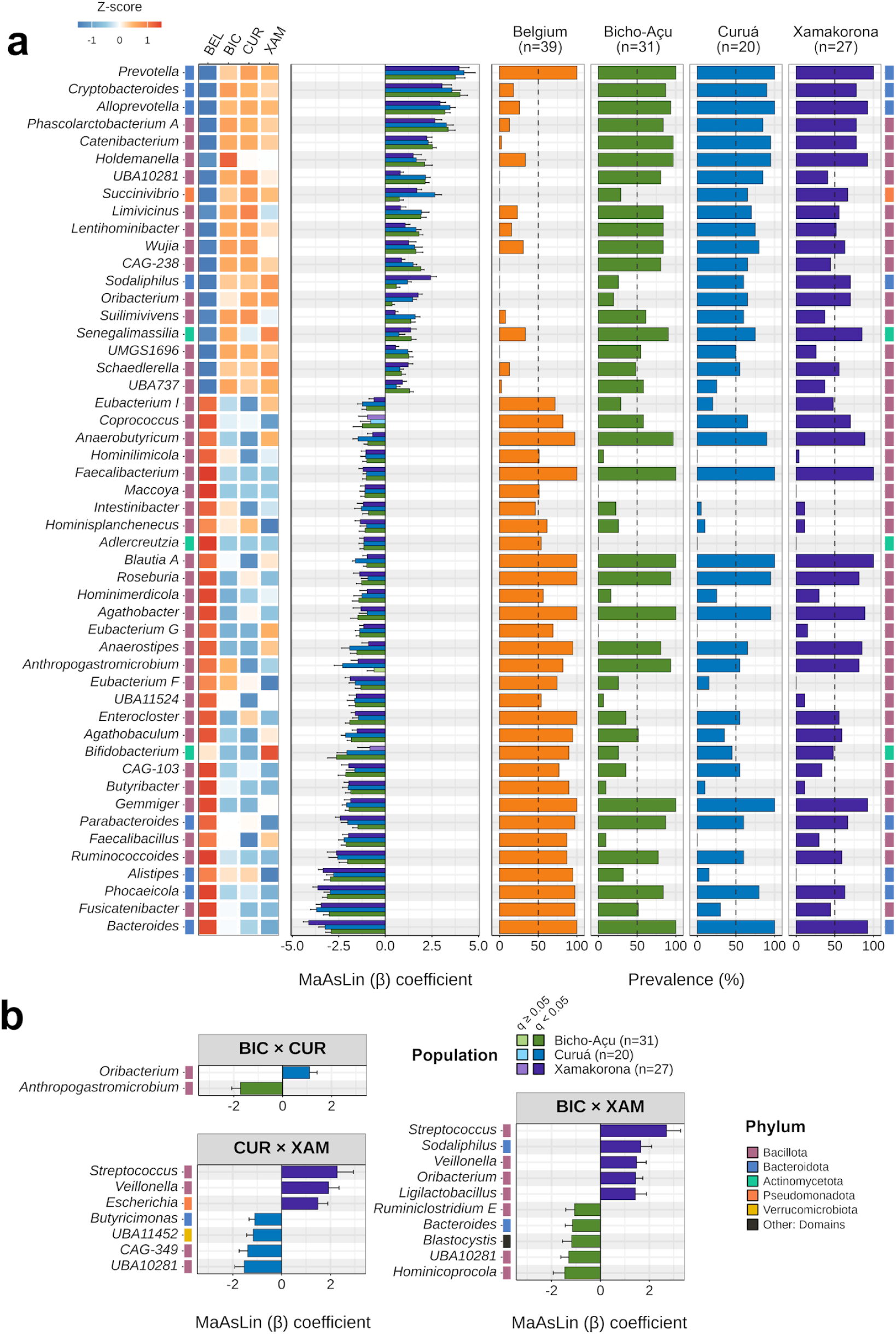
Genus-level differential abundance reveals asymmetric taxonomic turnover between Belgian and Yanomami microbiomes. **a,** Top 50 genera showing the greatest overall differences between BEL and the Yanomami populations, ranked according to the mean MaAsLin2 regression coefficient across BEL–BIC, BEL–CUR, and BEL–XAM contrasts. The left heatmap shows population-level mean relative abundances after centered log-ratio transformation and feature-wise Z-scaling. The center plot shows MaAsLin2 regression coefficients (β) ± standard error for each Yanomami population relative to BEL; muted colors indicate non-significant associations (*q* > 0.05). The right plot shows genus prevalence within each population, with the dashed vertical line indicating 50% prevalence. Phylum annotations are indicated by color-coded side bars. b, Differentially abundant genera identified in pairwise comparisons among Yanomami subpopulations; all genera meeting the significance criteria are shown as MaAsLin2 coefficient estimates. Genus relative abundances were TSS-normalized for MaAsLin2 analysis, and associations were considered significant at |β| > 1 and Benjamini–Hochberg-adjusted *q* < 0.05.

Differential abundance was also detected among the Yanomami subpopulations (**Fig. 2b**). BIC and CUR differed only in two taxa, whereas ten genera differed between BIC and XAM and seven between CUR and XAM. Within these comparisons, *Bacteroides* was enriched in BIC relative to XAM but did not differ between BIC and CUR. Conversely, *Oribacterium* was enriched in CUR and XAM relative to BIC, consistent with its lower prevalence in BIC and absence from BEL. These opposing abundance patterns, together with the intermediate occupancy patterns described above, support BIC as a possible transitional taxonomic configuration, combining selected urban-associated features with a broadly Yanomami-associated microbiome. These contrasts, nevertheless, remained substantially smaller than the differences that distinguished Yanomami subpopulations from BEL.

### Microbial association networks show an ordered change in connectivity

Genus-level co-occurrence networks differed substantially in their topology (**Fig. 3, Supplementary Table 5**). The number of nodes, each representing a genus retained in the corresponding population network, was slightly reduced in the urban cohort but remained relatively similar across the Yanomami subpopulations. However, the number of retained correlations, represented as edges connecting genera, increased more prominently in the sequence BEL (236), BIC (665), CUR (749), and XAM (1,423). The average number of connections per genus (mean node degree) followed the same ordering across populations. Targeted node-removal simulations showed corresponding differences in connectivity loss. After removing approximately 25% of nodes in descending order of degree, the normalized connectivity-loss metric was 95.4% in BEL, 75.79% in BIC, 49.1% in CUR, and 45.1% in XAM. The number of connector taxa, those exhibiting relatively high connectivity across network modules and therefore potentially linking distinct groups of genera, also increased from six in BEL to seven in BIC, eight in CUR, and 36 in XAM. Moreover, *Lachnospira* in XAM was the only taxon classified as a modular hub—a node with high connectivity and centrality—across all four networks (**Fig. 3c,d** and **Supplementary Table 5**).

**Figure 3.**
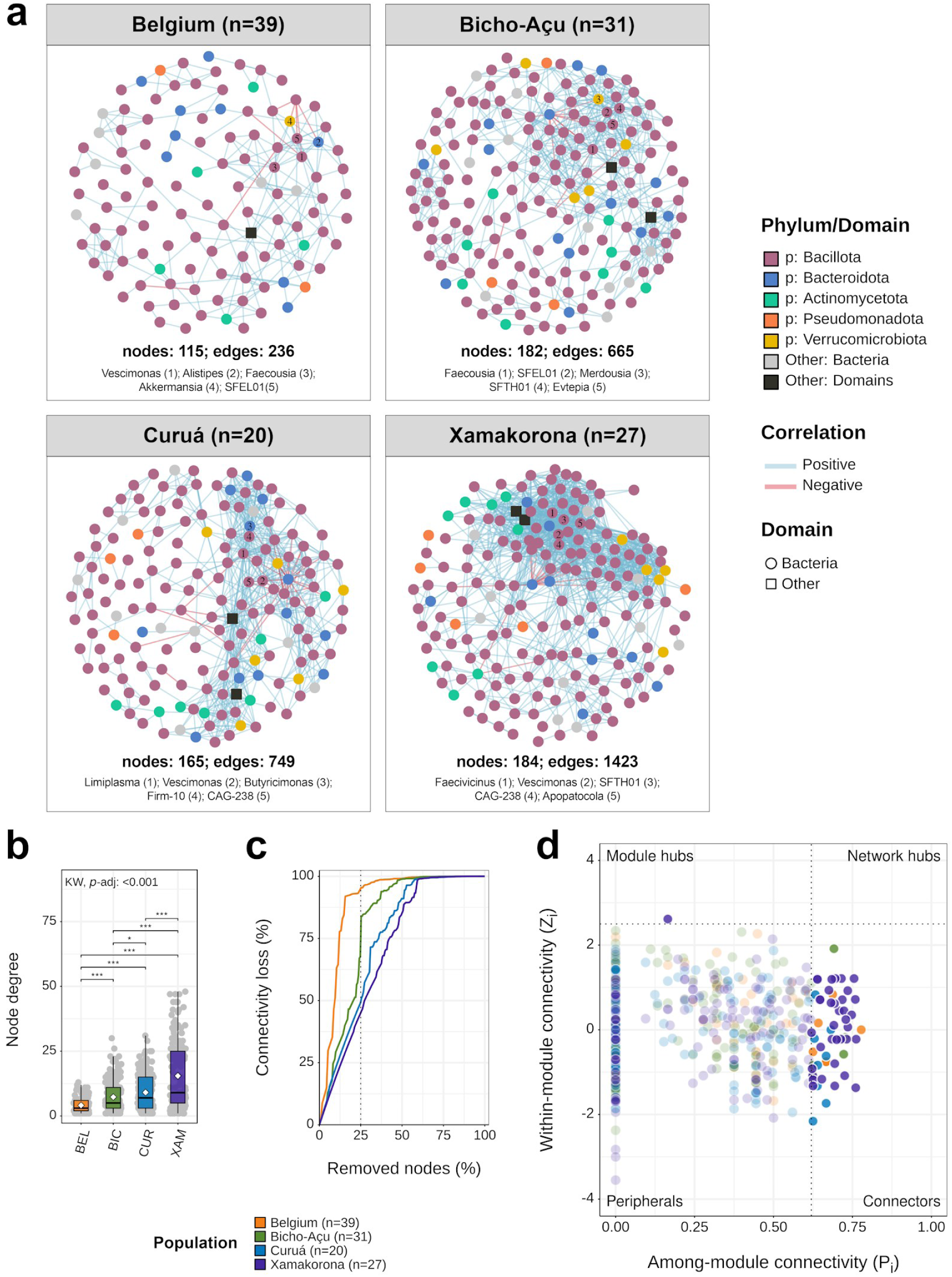
Genus-level co-occurrence networks show progressively reduced connectivity toward the urbanized population. Networks were constructed independently for each population. a, Genus-level co-occurrence networks based on Pearson correlations. Edges represent associations with |*r*| ≥ 0.5 and FDR-adjusted *P* ≤ 0.05; positive and negative correlations are distinguished by edge color. The five genera with the highest Kleinberg hub scores in each population are indicated. b, Mean node degree, representing the number of connections per genus. Mean ± SD node degree was 4.10 ± 2.89 in BEL, 7.31 ± 5.99 in BIC, 9.08 ± 7.07 in CUR, and 15.47 ± 12.81 in XAM. Node-degree distributions differed significantly among populations (Kruskal–Wallis *H* = 94.25, FDR-adjusted *P* <0.001), followed by pairwise two-sided Wilcoxon rank-sum tests with Benjamini–Hochberg FDR correction. c, Model-based network robustness during sequential removal of nodes in descending order of degree, shown as the progressive loss of network connectivity. The removal of 25% of the nodes is indicated by the dotted line. d, Zi–Pi plots showing within-module connectivity (Zi) and among-module connectivity (Pi) used to classify topological node roles. Thresholds were Zi = 2.5 and Pi = 0.62 (dotted lines), and modules were identified using the Louvain community detection algorithm. Exact network metrics and statistical results are provided in **Supplementary Table 5**. Significance for FDR-adjusted pairwise comparisons is indicated as \*\*\**P* < 0.001, \*\**P* < 0.01, \**P* < 0.05, and ∼*P* < 0.1.

Despite this divergence, several genera recurred among the five highest hub-centrality scores: *Vescimonas* in BEL, CUR, and XAM; *Faecousia* and *SFEL01* in BEL and BIC; *SFTH01* in BIC and XAM; and *CAG-238* in CUR and XAM, suggesting that some members may maintain their centrality roles across populations spanning a wide range of urbanization levels. Taken together, the ordered differences in connectivity, connector abundance, and simulated connectivity loss support a reorganization of microbial ecological relationships across the urbanization gradient. XAM exhibited the most densely connected association network, whereas BIC occupied an intermediate position between BEL and the less urban-influenced Yanomami subpopulations. This pattern is consistent with a less interconnected microbial community in populations with greater urban influence.

### Populations show extensive differentiation in metabolic potential

We next examined the functional potential encoded by these microbiomes through gene prediction and functional annotation (**Supplementary Table 3**). Profiling of the complete KEGG ortholog (KO) repertoire identified 5,906 KOs prevalent in at least 50% of individuals across all four populations and comparatively few population-exclusive KOs: 89 in BEL, 61 in BIC, 67 in CUR and 99 in XAM (**Fig. 4a**). KO Shannon diversity differed only modestly in magnitude but increased in population order, with XAM also differing from BIC (**Fig. 4b**). However, population-level differences in KO composition were more pronounced, accounting for 24% of Bray–Curtis variation (**Fig. 4c**). BEL was most strongly separated from each Yanomami subpopulation (pairwise *F* = 17.33–24.80, *P* < 0.001), although all three within-Yanomami comparisons were also significant (pairwise *F* = 3.34–6.09, *P* ≤ 0.003; **Supplementary Table 6**).

**Figure 4.**
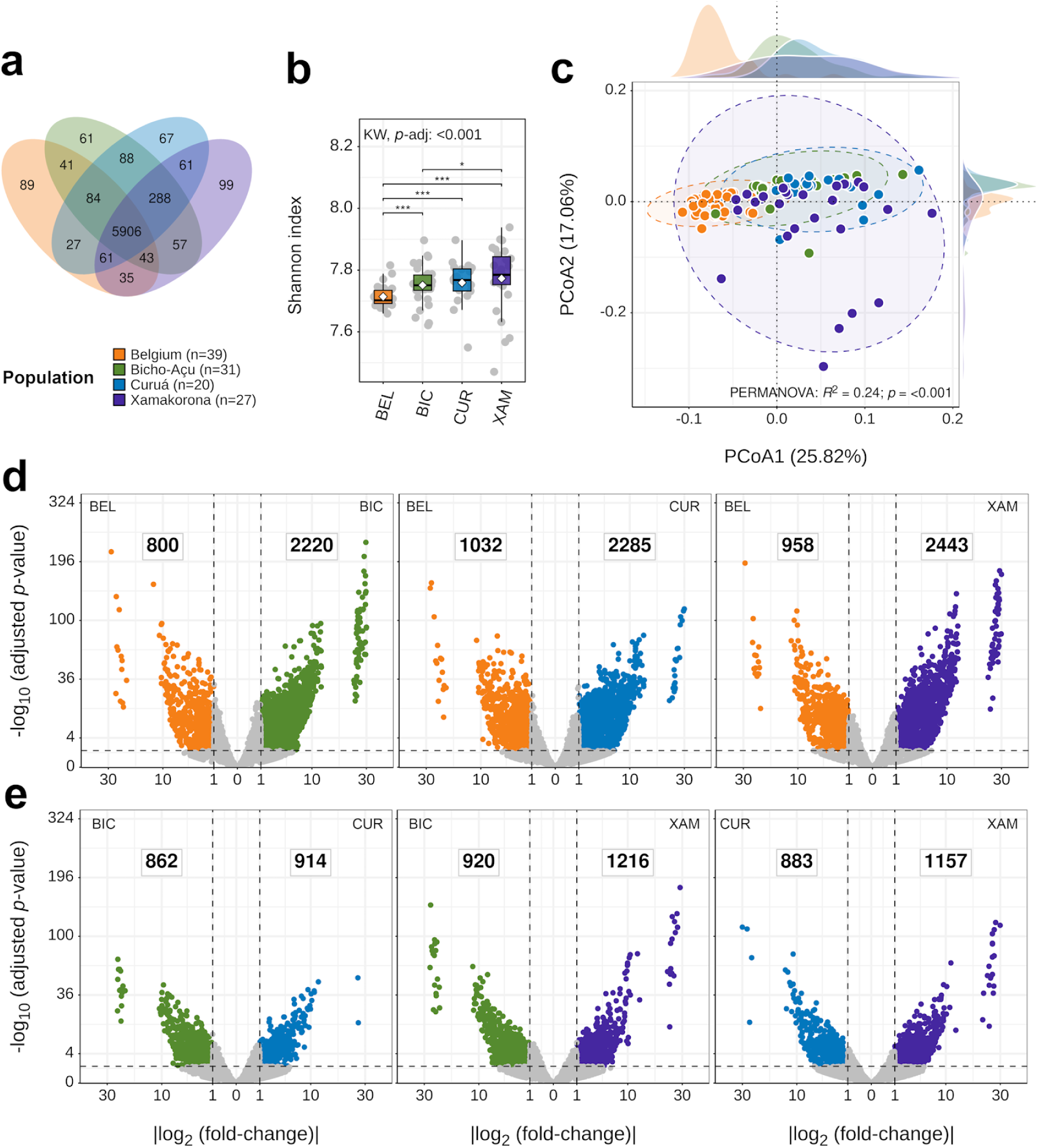
KEGG ortholog profiles reveal greater functional diversity and progressive functional divergence across the urbanization gradient. **a,** Venn diagram showing KOs shared among or exclusive to populations; KOs were considered present when detected in ≥50% of individuals within a population. b, Shannon diversity of KO profiles. Mean ± SD: BEL, 7.71 ± 0.03; BIC, 7.75 ± 0.06; CUR, 7.76 ± 0.07; XAM, 7.77 ± 0.11. Overall differences were assessed by Kruskal–Wallis test (*H* = 23.94, FDR-adjusted *P* < 0.001), followed by pairwise two-sided Wilcoxon rank-sum tests with Benjamini–Hochberg correction. c, PCoA of Bray–Curtis dissimilarities calculated from KO profiles. Density plots show the distribution of sample scores along PCoA1 (top) and PCoA2 (right), grouped according to the color coding in each panel. Population explained 24% of functional compositional variation (PERMANOVA: R2 = 0.24, *F* = 12.09, *P* < 0.001; 999 permutations). Pairwise PERMANOVA results are provided in **Supplementary Table 6. d**, Differentially abundant KOs between BEL and each Yanomami population, identified using DESeq2. Positive and negative log2 fold-change values indicate enrichment in Yanomami and BEL samples, respectively. e, DESeq2 differential-abundance comparisons among BIC, CUR, and XAM. In d,e, differential abundance was defined as |log2 fold-change| > 1 and FDR-adjusted *P* < 0.05; non-significant KOs are shown in gray. The number of differentially abundant KOs in the population are indicated on the corresponding side of each comparison.Square-root-scaled log2 fold-change axes are used to improve visualization of extreme values. Significance for FDR-adjusted comparisons is indicated as \*\*\**P* < 0.001, \*\**P* < 0.01, \**P* < 0.05, and ∼*P* < 0.1.

Despite the large functional core, differentiation extended across thousands of individual KOs (**Figs. 4d,e**). The BEL–BIC, BEL–CUR, and BEL–XAM comparisons yielded 3,020, 3,317, and 3,401 differentially abundant KOs, respectively. In each comparison, more KOs were enriched in the Yanomami population than in BEL. Differentiation within the Yanomami cohort was also extensive, comprising 1,776 KOs between BIC and CUR, 2,136 between BIC and XAM, and 2,040 between CUR and XAM. These differences reveal extensive redistribution of encoded metabolic potential within a largely shared KO repertoire. The divergence in KO profiles relative to BEL increased progressively from BIC to CUR and XAM, while the larger internal Yanomami contrasts also involved XAM. Together with the overall functional composition, this ordering supports BIC as a possible transitional configuration, retaining a broadly Yanomami-associated functional profile while showing the fewest KO-level abundance differences from the urban cohort.

Over-representation analysis placed these KO-level differences into KEGG Metabolism subclasses while testing enrichment specifically among differentially abundant KOs (**Fig. 5a**, **Supplementary Fig. 2a**). Metabolism of terpenoids and polyketides was the only subclass consistently over-represented among KOs enriched in the more urbanized member of every pairwise comparison (fold enrichment = 1.61–2.17, FDR-adjusted *P* = 3.73 × 10^−7^ to 0.026), as this function was consistently enriched in BEL relative to each Yanomami subpopulation, but also in BIC relative to CUR and XAM, and CUR relative to XAM. Xenobiotics biodegradation and metabolism provided the only additional significant subclass result and was over-represented among BIC-enriched KOs relative to CUR (fold enrichment = 1.51, FDR-adjusted *P* = 0.016).

**Figure 5.**
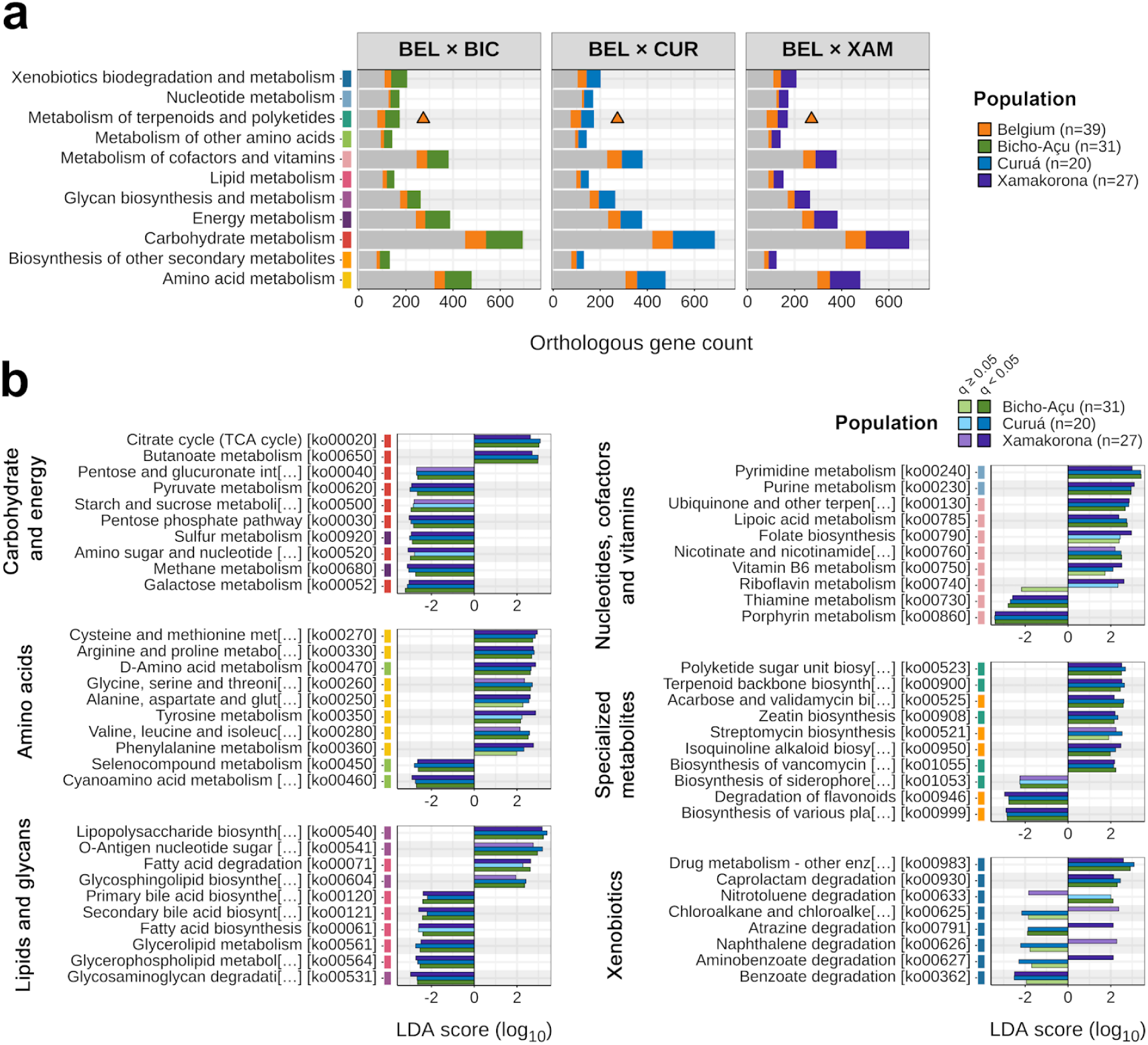
Urbanization is associated with shifts in the metabolic gene repertoire and pathway-level functional potential. **a,** Assignment of differentially abundant KOs identified by DESeq2 in BEL–BIC, BEL–CUR, and BEL–XAM comparisons to functional subclasses within the KEGG Metabolism class. Colored bars represent differentially abundant KOs assigned to each subclass, whereas gray bars represent non-differentially abundant KOs. Triangles denote KEGG metabolism subclasses significantly enriched among differentially abundant KOs by over-representation analysis using a hypergeometric test with Benjamini–Hochberg-adjusted *P* ≤ 0.05. Individual KOs were considered differentially abundant at |log2 fold-change| > 1 and FDR-adjusted *P* < 0.05. b, Top 10 KEGG metabolic pathways distinguishing BEL from each Yanomami population, identified by LEfSe, organized into functional groups. Pathways were retained at LDA effect size > 2, and Benjamini–Hochberg-adjusted *P* < 0.05; muted colors denote pathways not meeting the adjusted-*P* significance criterion. The annotation bar adjacent to pathway names indicates the KEGG metabolism subclass grouping used in a. Corresponding comparisons among Yanomami populations are shown in Supplementary Figure 2. Significance, where indicated, follows ***FDR-adjusted *P* < 0.001, \*\**P* < 0.01, \**P* < 0.05, and ∼*P* < 0.1.

LEfSe analysis of the relative abundance of summed KO counts assigned to each KEGG Metabolism pathway revealed broader functional differences (**Fig. 5b**). 60, 64, and 67 pathways distinguished BEL from BIC, CUR, and XAM, respectively (LDA > 2 and FDR-adjusted P < 0.05). Among pathways significant in all three comparisons, the largest recurrent BEL-associated effects were for porphyrin (LDA = 3.39–3.42), galactose (3.04–3.22), sulfur (2.89–3.02), and methane metabolism (2.73–3.12). The strongest recurrent Yanomami-associated effects involved lipopolysaccharide biosynthesis (LDA = 3.17–3.40), pyrimidine (2.99–3.42) and purine metabolism (2.94–3.10); butanoate metabolism was also consistently and strongly enriched across the three Yanomami subpopulations (2.70–2.98). Within the Yanomami cohort, only eight pathways distinguished BIC from CUR, compared with 38 between BIC and XAM and 28 between CUR and XAM (**Supplementary Fig. 2b**). This general asymmetry resembles the pattern observed in taxonomic marker assessment, where the greatest internal contrasts involve the XAM subpopulation, whereas the differences between BIC and CUR are comparatively smaller. This suggests possible selective changes, in which Yanomami populations under greater urban influence exhibit divergences in encoded functional potential relative to the XAM configuration.

Peptidoglycan biosynthesis and glutathione metabolism were among the strongest pathways recurrently enriched in XAM relative to both BIC and CUR (LDA = 3.09 and 3.04 for peptidoglycan biosynthesis, and 2.80 and 2.66 for glutathione metabolism, respectively). Conversely, glycosaminoglycan degradation and secondary bile acid biosynthesis, pathways involved in the microbial transformation of host-derived glycans and bile acids, were enriched in BIC and CUR relative to XAM (LDA = 2.61 and 2.61 for glycosaminoglycan degradation, and 2.14 and 2.34 for secondary bile acid biosynthesis, respectively). Both pathways were also enriched in BEL across all comparisons with Yanomami subpopulations. In the smaller BIC–CUR set, O-antigen nucleotide-sugar biosynthesis showed the largest effect and was enriched in CUR (LDA = 2.92, FDR-adjusted *P* = 0.048), whereas the BIC-associated pathways were dominated by xenobiotic-degradation functions, including benzoate degradation and cytochrome-P450-mediated xenobiotic metabolism.

### Butyrate biosynthetic potential varies by pathway preference and taxonomic contributors

Butyrate is a major microbiota-derived short-chain fatty acid (SCFA), and butanoate metabolism was consistently enriched in all three Yanomami subpopulations at the KEGG pathway level; therefore, we examined whether this difference reflected specific biosynthetic routes. We reconstructed four main microbial butyrate biosynthesis routes, the acetyl-CoA (P1), 4-aminobutyrate/succinate (P2), glutamate (P3), and lysine (P4) pathways, together with terminal conversion reactions (terminal enzymes, T; **Fig. 6a**). At the route level, summed gene abundance did not differ significantly among populations for P1, P2, or T (**Fig. 6b**). Individual KOs within P1 and T nevertheless showed differential abundance, revealing redistribution across specific enzymatic steps despite similar aggregate abundances (**Fig. 6a** and **Supplementary Fig. 3**). By contrast, summed P3 gene abundance declined steadily from 599.45 counts per million (CPM) in BEL to 516.20 in BIC, 496.97 in CUR, and 428.05 in XAM (*H* = 18.71, FDR-adjusted *P* < 0.001). P4 was also lower in the Yanomami subpopulations, averaging 299.32 CPM in BEL, 203.40 in BIC, 229.93 in CUR, and 157.99 in XAM (*H* = 26.51, FDR-adjusted *P* < 0.001).

**Figure 6.**
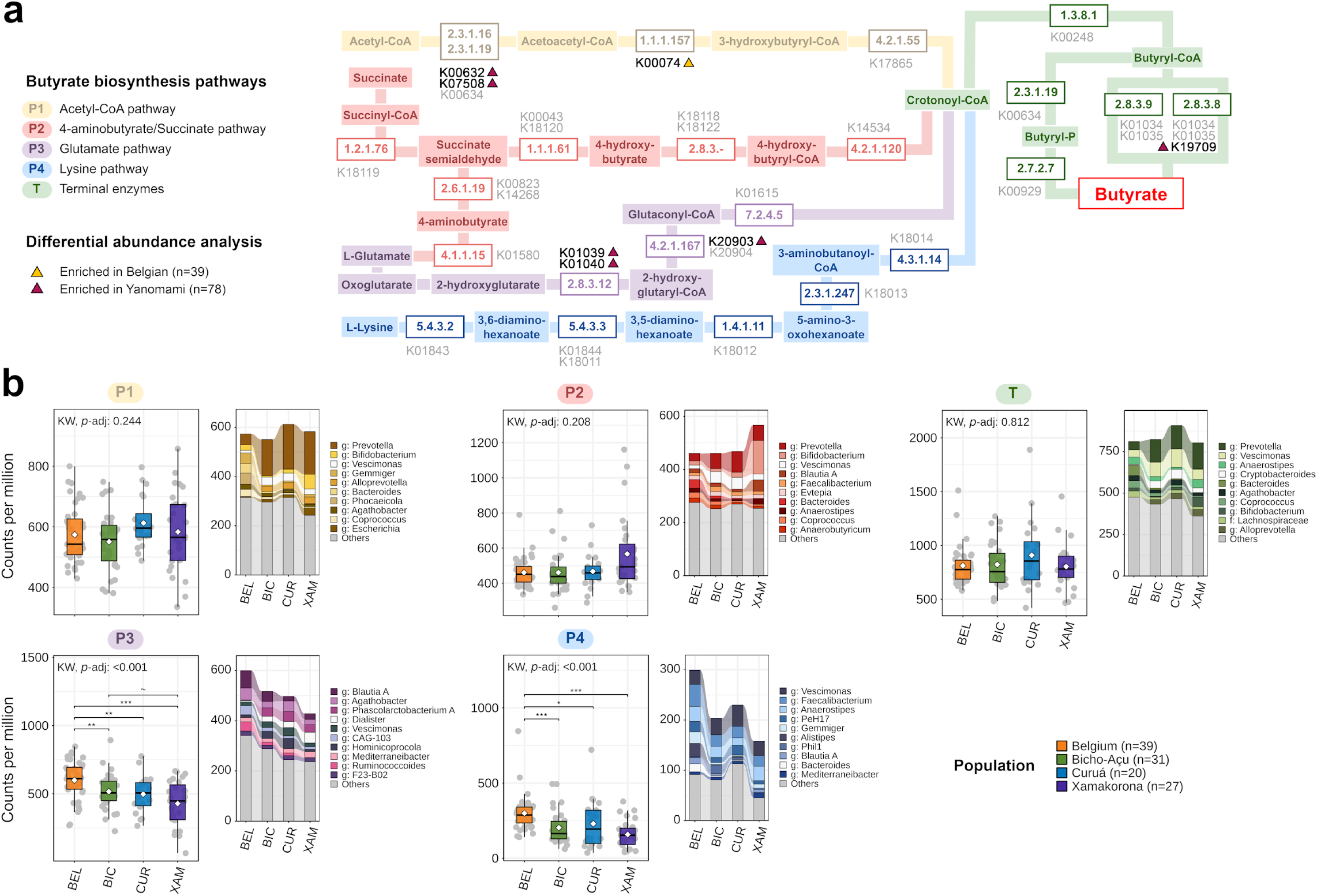
Microbial butyrate biosynthetic potential differs by pathway and taxonomic contribution across populations. **a,** Reconstruction of the four major microbial butyrate biosynthesis pathways based on Frolova et al. and Kisuse et al. as described in Methods. KOs detected in the metagenomic dataset were mapped to the corresponding enzymatic reaction steps according to their EC assignments. Differential abundance was evaluated by DESeq2 between BEL and pooled Yanomami samples (n = 78). KOs shown in gray were not significantly different, whereas bold black KOs met the differential-abundance criteria (|log2 fold-change| > 1 and FDR-adjusted *P* < 0.05); triangles indicate the population in which the KO was enriched. Population-specific pairwise comparisons are shown in **Supplementary Figure 3. b**, Total abundance of gene copies assigned to P1–P4 and T, expressed as counts per million (CPM), together with the mean taxonomic contribution of the 10 most abundant genus-level taxa within each pathway and population. Mean ± SD CPM values were: P1 = BEL, 573.90 ± 93.88; BIC, 550.72 ± 99.06; CUR, 612.59 ± 84.29; XAM, 583.26 ± 130.69; P2 = BEL, 460.58 ± 82.20; BIC, 460.67 ± 111.45; CUR, 467.35 ± 97.46; XAM, 566.84 ± 211.26; P3 = BEL, 599.45 ± 141.06; BIC, 516.20 ± 131.05; CUR, 496.97 ± 136.10; XAM, 428.05 ± 168.87; P4 = BEL, 299.32 ± 119.75; BIC, 203.40 ± 109.27; CUR, 229.93 ± 166.44; XAM, 157.99 ± 91.81; T = BEL, 811.16 ± 175.92; BIC, 822.72 ± 208.46; CUR, 908.25 ± 332.62; XAM, 803.39 ± 208.67. Overall differences were assessed by Kruskal–Wallis tests: P1, *H* = 4.70, FDR-adjusted *P* = 0.244; P2, *H* = 5.74, *P* = 0.208; P3, *H* = 18.71, *P* = 0.001; P4, *H* = 26.51, *P* < 0.001; T, *H* = 0.95, *P* = 0.812. Significant omnibus tests were followed by pairwise two-sided Wilcoxon rank-sum tests with Benjamini–Hochberg correction. Significance for FDR-adjusted pairwise comparisons is indicated as \*\*\**P* < 0.001, \*\**P* < 0.01, \**P* < 0.05, and ∼*P* < 0.1.

Analysis of taxonomic contributions showed that similar aggregate pathway abundances could be encoded by different community members (**Fig. 6b**). In P1, *Prevotella* accounted for 7.5% of the mean encoded abundance in BEL but 26.5% in BIC and 29.8% in both CUR and XAM. By contrast, *Phocaeicola* and *Bacteroides* contributed 7.9% and 6.9%, respectively, in BEL but only 0.1–0.2% and 0.4–1.1% across the Yanomami cohorts. *Prevotella* similarly contributed 6.1% of terminal-reaction abundance in BEL and 15.7–19.8% across the Yanomami microbial communities, accompanied by greater contributions from *Vescimonas* and *Cryptobacteroides* in BIC and CUR. P2 in XAM had a distinct configuration in which *Bifidobacterium* contributed 22.2% of the encoded pathway abundance. For P3, the decline in total abundance was accompanied by a shift from *Blautia A* and *Agathobacter*, the two leading named contributors in BEL, to *Dialister* and *Phascolarctobacterium A* in XAM. P4 was supported principally by *Faecalibacterium*, *Anaerostipes,* and *Vescimonas*, but their relative contributions varied across populations.

Individual KO-level differences further revealed redistribution of abundance across specific enzymatic steps. Relative to BEL, pooled Yanomami samples were enriched in six butyrate-associated KOs: two assigned to P1, three to P3, and one to terminal conversion reactions (log_2_ fold-change = 1.51–4.48; all FDR-adjusted *P* < 0.001; **Fig. 6a**). By contrast, the P1-associated KO K00074 was enriched in BEL (log_2_ fold-change = −1.38; FDR-adjusted *P* = 2.55 × 10^−16^). Population-specific comparisons generally recapitulated these patterns (**Supplementary Fig. 3**). Within P1, K00074 was more abundant in BEL than in each Yanomami subpopulation (Yanomami-versus-BEL log_2_ fold-change = −2.05 to −0.86; all FDR-adjusted *P* < 0.001), whereas K00632 was more abundant in all three Yanomami subpopulations than in BEL (log_2_ fold-change = 1.60–2.58; FDR-adjusted *P* < 0.001). K00632 was also higher in CUR and XAM than in BIC, although with comparatively modest effect sizes (log_2_ fold-change = 0.99 and 0.89; FDR-adjusted *P* = 0.027 and 0.030, respectively). K07508 was enriched in BIC and XAM relative to BEL.

Despite the overall higher aggregate abundance of P3 in BEL, three KOs of this biosynthetic pathway were enriched in Yanomami subpopulations relative to the urban cohort (log2 fold-change = 1.27–4.14; FDR-adjusted *P* <0.001). However, their CPM abundance profiles tended to be higher in BIC and CUR and lower in BEL and XAM. Finally, a terminal conversion-associated KO, K19709, showed the clearest ordered pattern, with enrichment of CUR and XAM relative to BEL, of CUR relative to BIC, and of XAM relative to both BIC and CUR (log_2_ fold-change = 1.25–4.03; all FDR-adjusted *P* ≤ 0.035). Together, these patterns distinguish the aggregate abundance of pathways from the individual redistribution of specific P1, P3, and terminal steps, including routes whose total abundance remained unchanged. Therefore, they indicate route-specific reorganization of encoded butyrate biosynthetic potential.

### CAZyme profiles distinguish populations and suggest greater non-starch polysaccharide-processing potential in the more urbanized cohort

Carbohydrate-active enzymes (CAZymes) mediate the breakdown, biosynthesis and modification of glycans, thereby shaping how gut microbial communities access dietary and host-derived carbohydrates. CAZyme profiles contained a core of 353 families prevalent across all four populations, with 19 families exclusive to BEL, five to BIC, six to CUR and seven to XAM at the 50% prevalence threshold (**Fig. 7a**). CAZyme Shannon diversity differed among populations (Kruskal–Wallis *H* = 43.31, FDR-adjusted *P* < 0.001), with means of 4.65 in BEL, 4.60 in BIC, 4.63 in CUR and 4.47 in XAM (**Fig. 7b**). Population explained 22% of Bray–Curtis variation in CAZyme composition (PERMANOVA: R^2^ = 0.22, *F* = 10.67, *P* < 0.001; **Fig. 7c**). All pairwise comparisons were significant, with larger effects in BEL–Yanomami contrasts (pairwise *F* = 13.61–18.47, *P* < 0.001) than among Yanomami subpopulations (pairwise *F* = 2.20–6.82; **Supplementary Table 7**).

**Figure 7.**
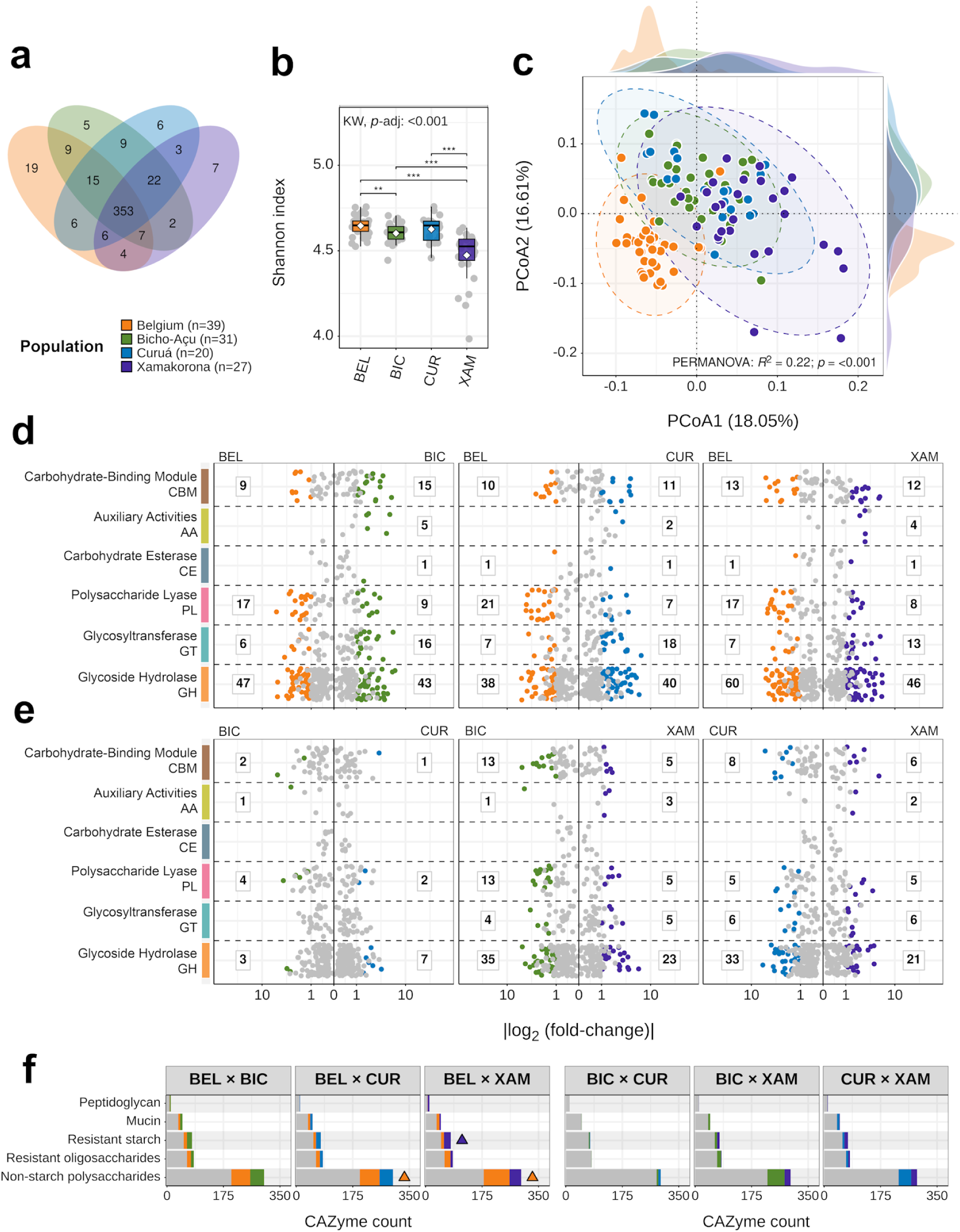
Carbohydrate-active enzyme repertoires reveal population-specific differences in carbohydrate-processing potential. **a,** Venn diagram showing CAZymes exclusive or shared among populations; CAZymes were considered present when detected in ≥50% of individuals within a population. b, Shannon diversity of CAZyme profiles. Mean ± SD: BEL, 4.65 ± 0.06; BIC, 4.60 ± 0.05; CUR, 4.63 ± 0.08; XAM, 4.47 ± 0.15. Overall differences were significant (Kruskal–Wallis *H* = 43.31, FDR-adjusted *P* < 0.001), followed by pairwise two-sided Wilcoxon rank-sum tests with Benjamini–Hochberg correction. c, PCoA of Bray–Curtis dissimilarities calculated from CAZyme profiles. Density plots show the distribution of sample scores along PCoA1 (top) and PCoA2 (right), grouped according to the color coding in each panel. Population explained 22% of compositional variation (PERMANOVA: R2 = 0.22, *F* = 10.67, *P* < 0.001; 999 permutations). Pairwise PERMANOVA results are provided in **Supplementary Table 7. d**, Differential abundance of CAZymes between BEL and each Yanomami population identified using DESeq2. Positive and negative log2 fold-change values indicate enrichment in Yanomami and BEL samples, respectively. e, Corresponding differential-abundance comparisons among BIC, CUR, and XAM. In d,e, significance was defined as |log2 fold-change| > 1 and FDR-adjusted *P* < 0.05; non-significant CAZymes are shown in gray. The number of differentially abundant CAZymes in the population is indicated on the corresponding side of each comparison. Square-root-scaled log2 fold-change axis is used to improve visualization of extreme values. f, Assignment of differentially abundant CAZymes to Cayman level-2 substrate categories. The left set shows BEL–BIC, BEL–CUR, and BEL–XAM comparisons, and the right set shows comparisons among Yanomami populations. Colored bars indicate differentially abundant CAZymes assigned to substrate categories, whereas gray bars indicate non-differentially abundant CAZymes. Triangles indicate substrate categories enriched among differentially abundant CAZymes by over-representation analysis using a hypergeometric test with Benjamini–Hochberg-adjusted *P* ≤ 0.05. Significance for FDR-adjusted comparisons is indicated as \*\*\**P* < 0.001, \*\**P* < 0.01, \**P* < 0.05, and ∼*P* < 0.1.

DESeq2 identified 168, 156 and 182 differentially abundant CAZymes in BEL–BIC, BEL–CUR and BEL–XAM comparisons, respectively (**Fig. 7d**). These features were distributed relatively evenly by direction: 79 were enriched in BEL and 89 in BIC; 77 in BEL and 79 in CUR; and 98 in BEL and 84 in XAM. Glycoside hydrolases (GHs) accounted for the largest share of differences across all contrasts, but other enzyme classes showed directional asymmetry. Polysaccharide lyases (PLs) were consistently more numerous among BEL-enriched than Yanomami-enriched features, whereas glycosyltransferases (GTs) showed the opposite pattern. Differences among Yanomami subpopulations were most limited between BIC and CUR, with 20 differentially abundant CAZymes, and was greater between BIC and XAM (107) and between CUR and XAM (92; **Fig. 7e**). The BIC–XAM contrast also contained a marked PLs imbalance, with 13 features enriched in BIC and five in XAM, reproducing the BEL versus Yanomami pattern.

Over-representation analysis tested which substrate annotations were concentrated among differentially abundant CAZyme families (**Fig. 7f**). Non-starch-polysaccharide (NSP) functions were over-represented among BEL-enriched CAZymes relative to CUR (fold enrichment = 1.18, FDR-adjusted *P* = 0.026) and XAM (fold enrichment = 1.16, FDR-adjusted *P* = 0.019). Resistant-starch functions were over-represented among XAM-enriched CAZymes relative to BEL (fold enrichment = 1.68, FDR-adjusted *P* = 0.021). No substrate class showed enrichment in the within-Yanomami population comparisons. The genera encoding these CAZyme profiles also differed consistently between BEL and the Yanomami microbial communities (**Supplementary Fig. 4a**). In BEL, *Bacteroides* was the leading named contributor to mucin, NSP, resistant-oligosaccharide and peptidoglycan categories, accounting for 9.0–14.1% of their mean encoded abundances, while *Ruminococcoides* was the leading contributor to resistant starch (12.4%). In the Yanomami population, *Prevotella* was the leading contributor in all five categories. *Cryptobacteroides* contributed prominently in BIC and CUR, particularly to mucin, NSP and resistant oligosaccharides, whereas *Bifidobacterium* made a larger contribution in XAM, including 17.9% of resistant-starch and 12.0% of NSP potential.

Total CAZyme abundance assigned to individual substrates revealed several distinct configurations (**Supplementary Fig. 4b**). Xylan-targeting CAZymes were most abundant in BEL (7.54%, versus 6.60–6.80% in the Yanomami cohorts), as were those assigned to arabinoxylan (6.45%, versus 5.42–5.76%), arabinan (5.76%, versus 4.68–4.84%) and arabinogalactan (5.72%, versus 4.47–4.76%; all omnibus *P* < 0.001), all NSPs. Starch-targeting CAZymes were instead most abundant in XAM (6.24%, versus 5.17–5.80%; *H* = 25.45, *P* < 0.001), whereas mucin-glycan CAZymes were highest in BIC (6.92%, versus 6.19% in BEL, 6.62% in CUR and 6.38% in XAM; *H* = 25.57, *P* < 0.001). Thus, population differences in carbohydrate-processing potential encompass changes in the abundance and substrate-preference distribution of CAZyme families, as well as the replacement of the main taxa carrying those functions.

## Discussion

The gut microbiomes of Indigenous populations remain underrepresented in global metagenomic datasets, limiting our understanding of the ecological diversity on which urbanization acts^21^. Addressing this gap is increasingly urgent as Indigenous peoples, including Yanomami communities, experience growing exposure to urban-associated influences^20,24^. Although differences in gut microbiome composition between populations at opposite ends of the urbanization spectrum are well documented, how these differences initially emerge and at what levels of urban exposure they become detectable remain incompletely understood. By profiling a cohort of 78 Yanomami participants from three distinct subpopulations, this study moves beyond comparisons of lifestyle endpoints to examine urban-associated changes within a shared population context. The resulting picture is that urbanization-associated differences remodel distinct ecological layers at different rates: the abundance and population prevalence of taxa, the organization of microbial associations, the relative genomic investment in metabolic processes, and the taxa and enzymatic routes through which substrates may be transformed. This layered response extends evidence that microbiome transitions are strongly contingent on geography, culture, and population history, even within the same broad process of urbanization^14–17,25^.

The taxonomic results suggest two complementary modes of remodeling. The opposing abundances of nearly ubiquitous *Prevotella* and *Bacteroides* indicate a change in the competitive weighting of organisms that remain widespread, whereas the contrasting prevalence profiles of other genera indicate differences in population-level persistence or occupancy. These processes are ecologically distinct, as changes in abundance may reflect altered substrate supply, competition, or growth efficiency within an established microbial community; changes in occupancy may additionally involve transmission, colonization thresholds, environmental reservoirs, or the ability to recover after perturbation^26,27^. Urbanization can therefore reorganize a microbiome without immediately eliminating its dominant lineages, while other, less ubiquitous organisms may be progressively lost from the population-level microbial pool^9,11,25^. Recent evidence for the selective disappearance of cellulose-degrading guilds provides a particularly clear example of the latter process^13^. At the same time, the non-monotonic alpha-diversity pattern in our study argues against using richness or Shannon diversity as a sufficient measure of urbanization-associated disruption. BIC was the most diverse population, while BEL and XAM had similar Shannon diversity despite occupying opposite ends of our urbanization spectrum. Specific taxa abundance, prevalence, and metabolic potential may therefore change before or independently of any reduction in overall diversity.

The internal Yanomami comparisons further suggest that the order in which microbiome dimensions respond may be as informative as the final endpoint differences. BIC retained the dominant Yanomami taxonomic configuration while showing selected features oriented toward the more urban-associated state, including greater representation of *Bacteroides*, host-glycan- and bile-related functions relative to XAM, as well as an intermediate network configuration between the urban BEL and the less urbanized CUR and XAM. We therefore interpret BIC as a possible transitional mosaic on specific ecological axes. Human dietary interventions demonstrate that altered substrate supply might rapidly change microbial abundance and gene expression^10^. Longitudinal observations following migration to the United States likewise show increasing *Bacteroides*-to-*Prevotella* ratios within months, even while strain profiles remain largely stable^28^. These findings provide a biological basis for interpreting BIC as a configuration in which changing resource availability and competitive relationships favor selected urban-associated traits within a broadly retained Yanomami-associated community. Comparable combinations of retained non-urbanized and urban-associated features have been described in other Amazonian and South African populations^14,16,29^. BIC therefore provides a compelling setting to test whether shifts in microbial abundance, substrate-use potential and association structure precede broader community turnover. Further longitudinal sampling would be useful to establish the direction and persistence of urbanized-associated microbial features.

Genus co-occurrence network topology offers a systems-level view of this reorganization. The ordered increase in connectivity from BEL through BIC and CUR to XAM reveals increasingly extensive associations among microbial abundances in the less urbanized populations. One plausible explanation is a more interconnected trophic architecture, in which primary polysaccharide degraders release oligosaccharides or fermentation products used by secondary consumers. *Prevotella*-centered association networks have been described in non-industrialized settings, and experimental work demonstrates how polysaccharide breakdown can support other community members^30–32^. A complementary explanation is that multiple taxa respond independently to the same dietary or host conditions. Differences in these shared conditions among participants could therefore reshape network connectivity through coordinated responses to resource availability, as supported by models of habitat-dependent covariation in the human microbiota^33^. These mechanisms may coexist, as urbanization may reorganize both metabolic dependencies among taxa and the environmental conditions to which they respond. Their ecological consequences depend on the underlying relationships, because trophic exchange can facilitate resource use while also transmitting perturbations between dependent partners; theory shows that strong positive feedback among cooperating taxa can destabilize communities^34^. The networks here cannot distinguish these mechanisms, and their simulated robustness remains a topological property rather than a measurement of *in vivo* resilience. Nevertheless, their unusually coherent population ordering generates a testable hypothesis about whether urbanization may simplify or reorganize the trophic dependencies that link microbial guilds.

The populations shared much of their KO repertoire yet differed extensively in the relative abundance of gene families, indicating that urbanization-associated functional differentiation largely involved redistribution of shared capacities. This reorganization extended beyond the BEL–Yanomami contrast: BIC and CUR differed in fewer KOs and metabolic pathways than either did from XAM, indicating substantial functional differentiation within the Yanomami populations. A more consistently ordered signal emerged for terpenoid and polyketide metabolism: KOs assigned to this subclass were over-represented among those enriched in the more urbanized population in every pairwise comparison. While interesting in terms of signaling regarding urban influence, the biochemical breadth of these KEGG subclasses limits their utility as evidence of a specific exposure or substrate. In any case, their enrichment suggests broad-level candidate mechanisms undergoing functional reorganization.

The host-glycan and bile-related findings provide a plausible example of how metabolic reallocation could accompany taxonomic turnover. Flexible glycan foragers can switch between dietary and host-derived carbohydrates, potentially maintaining growth when accessible plant substrates change. In mice, deprivation of microbiota-accessible dietary carbohydrates drives increased consumption of mucus glycans and erosion of the mucus barrier^35^, whereas metaproteomic evidence shows that dietary fibers and mucin glycans can also be used in parallel rather than as mutually exclusive resources^36^. The enrichment of glycosaminoglycan-degradation genes in BEL and in BIC and CUR relative to XAM is therefore compatible with expansion of host-glycan-adapted niches, potentially facilitated by the greater representation of *Bacteroides* and other organisms with flexible glycan-utilization systems^37^. Glycosaminoglycans and mucin glycans are chemically distinct, however, so this should be treated as a broader host-glycan usage hypothesis rather than evidence of increased potential for mucus degradation. In this sense, a parallel gnotobiotic high-fat-diet experiment using the same donor microbiotas did not detect differences in mucus-layer thickness between mice receiving Belgian and Yanomami microbial communities (Thorp et al., *in prep.*).

The secondary bile acid metabolism signal raises a related ecological possibility. Bile acids simultaneously provide substrates for specialized bacterial metabolism, impose ecological selection through their detergent and antimicrobial properties, and participate in enterohepatic signaling. Supplementation with a 7α-dehydroxylating bacterium can modify colonization resistance through secondary-bile-acid-associated effects^38^; microbiota-dependent changes in bile-acid composition regulate intestinal FXR signaling and hepatic bile-acid synthesis^39^; and secondary-bile-acid depletion can promote intestinal inflammation, partly through reduced TGR5 signaling^40^. Conversely, excessive or sustained exposure to the hydrophobic secondary bile acids deoxycholic acid and lithocholic acid can disrupt cellular membranes, induce oxidative stress, and promote damage to DNA, mitochondria, and endoplasmic reticulum^41^. Bile-acid dysregulation might also impair tight-junction integrity, epithelial renewal, Paneth- and goblet-cell function, and antimicrobial defense, while selecting for bile-tolerant or opportunistic microorganisms^42^. Thus, altered representation of microbial secondary bile-acid pathways could have beneficial or detrimental consequences. Thorp et al. (*in prep*), through a gnotobiotic experiment, showed that secondary bile acid biosynthesis was enriched in mice receiving microbial communities of Belgian origin following the high-fat-diet challenge, reproducing the direction observed in the metagenomic contrast seen here. Also, recipients of the Yanomami microbiota exhibited an enrichment of transcripts related to hepatic bile acid metabolism, as well as reduced weight gain and alterations in hepatic lipid storage.

The consistent enrichment of the broad KEGG butanoate metabolism pathway in the Yanomami subpopulations prompted us to examine whether this signal could be determined into differences among specific butyrate biosynthetic routes. However, such enrichment is not uniform across all route-associated gene abundances. Summed abundances of P1, P2, and the terminal reactions did not differ significantly among populations, whereas P3- and P4-associated genes were more strongly represented toward BEL. The acetyl-CoA route (P1) is predominantly associated with carbohydrate-derived carbon, whereas the glutamate-derived (P3) and lysine (P4) routes draw on amino-acid-related substrates^43–45^. The greater summed representation of P3- and P4-associated genes in BEL, and partially with BIC (in relation to XAM), is therefore compatible with a greater genetic representation of amino-acid-associated routes of butyrate biosynthesis, particularly through lysine-associated metabolism. However, the divergent behavior of certain individual P3-KOs is particularly noteworthy, given that some enzymatic steps were more strongly represented in Yanomami microbiomes.

Similarly, although the combined abundance of KOs associated with terminal enzymes did not differ across populations, K19709 (Acetate CoA-transferase; involved in the final conversion of butyrate) exhibited a progressive enrichment towards the less urbanized communities. A consistent pattern at the broader level of SCFA-associated metabolism was observed in the gnotobiotic high-fat-diet assays (Thorp et al., *in prep.*), where Yanomami microbial communities showed enrichment of propanoate and central-carbon metabolic pathways alongside reduced metabolic dysfunction. Together, these findings suggest that urbanization is associated with a broader reorganization of encoded SCFA metabolism. As SCFAs participate in the regulation of epithelial barrier integrity, immune homeostasis, and host energy metabolism, shifts in the pathways might have important consequences for intestinal and metabolic physiology^46^.

Comparative studies have often reported enrichment of fiber-targeting CAZymes in non-industrialized microbiomes and loss of specialized cellulose-degrading organisms with industrialization^13,47^. Our findings refine this picture, whereas BEL showed stronger non-starch-polysaccharide-associated signals, and XAM showed resistant-starch-associated enrichment. Fiber, therefore, encompasses substrates that might favor different microbial strategies, making aggregate fiber-degradation capacity an insufficient description of these population differences. Our field observations and participant interviews indicated that processed foods were widely available frequently consumed in BIC and CUR. This pattern was not observed in XAM, where remoteness limited access to urban food supplies. Published syntheses of subsistence-based Amazonian diets describe reliance on starchy staples such as manioc or cassava and plantain, supplemented by a relatively restricted variety of other foods^48^, while research in other Yanomami communities documents frequent consumption of regional, *in natura* or minimally processed foods, but also substantial and strongly village-dependent exposure to ultra-processed foods^24^, illustrating that dietary transitions within Yanomami territory are heterogeneous. Differences in food sources and preparation methods may favor specialized carbohydrate-processing repertoires. For example, cooking, alters starch accessibility and other plant-food properties^49^, and structurally distinct resistant starches select different microorganisms and fermentation outputs^50^.

The shift of contributions from *Bacteroides* toward *Prevotella* and *Cryptobacteroides*, together with the prominent *Bifidobacterium* contribution to resistant-starch-associated genes in XAM, suggests changes in the organisms initiating carbohydrate breakdown and supplying metabolites to downstream consumers. Although *Bifidobacterium* is typically abundant in the infant gut^51^ and children represented a greater proportion of the XAM, this genus was also highly represented among adults in this population (data not shown). Nevertheless, experimental cocultures show that starch-utilizing *Bifidobacterium* species are able to support butyrate-producing partners through metabolic cross-feeding^52^. This provides a possible connection between XAM’s carbohydrate-processing signature and the organization of community fermentation. It also identifies a potential intervention target: matching carbohydrate structures to the organisms and metabolic partnerships present in a community. In a randomized crossover trial, resistant-starch supplementation improved body weight and insulin sensitivity in adults with overweight or obesity; accompanying experiments showed that *B. adolescentis* supplementation protected mice against diet-induced obesity^53^. In this sense, the XAM configuration associated with resistant starch could provide a useful basis for generating hypotheses about substrate-responsive microbial functions relevant to metabolic health that may be sensitive to urbanization.

## Conclusions

Together, our findings suggest that urbanization is associated with taxonomic turnover and a broader reorganization of microbial functions across the community. Importantly, the Yanomami subpopulations exhibited distinct microbiome configurations associated with differing degrees of urban influence. XAM, the least urban-exposed population, exhibited the taxonomic, network, and functional features that most distinguished the Yanomami microbiomes from the Belgian cohort. CUR, the intermediate subpopulation, showed specific differences from this configuration, whereas BIC, the most urban-exposed group, exhibited the strongest combination of characteristics associated with the urban environment within the broader Yanomami microbiome. Its taxonomic alterations, intermediate network structure, and specific shifts in metabolic potential suggest that the BIC may represent a transitional microbiome profile in which urban markers coexist with a predominantly Yanomami-associated signature. Longitudinal studies will be needed to determine whether this profile reflects an ongoing transition or a relatively stable state. Complementary results in animal models transplanted with these microbiotas lend physiological relevance to this process by demonstrating differences in susceptibility to diet-induced metabolic dysfunction. Linking the signatures identified here to the organisms, substrates and metabolites responsible for those effects could guide targeted microbiota interventions aimed at supporting metabolic health.

## Methods

### Ethical approval, participant recruitment, and sample collection

Procedures involving Yanomami participants were approved by the Brazilian Research Ethics Committee system (CEP/CONEP; no. 08069019.4.0000.5594). Recruitment of Belgian participants was approved by the Ethics Committee of Ghent University and Ghent University Hospital (project code ONZ-2023-0659). The participating Yanomami communities and their local leaders were consulted before the study and agreed to its implementation. Informed consent was obtained from all participants through procedures adapted to their cultural context, with the assistance of interpreters fluent in Yanomami languages and Portuguese. For participants younger than 18 years, consent was obtained from their parents or legal guardians. In accordance with guidelines for collaborative research and community engagement, preliminary findings were shared with members of the Yanomami communities using culturally appropriate approaches.

The study included 117 participants from four populations differing in geographic setting and exposure to urban-associated influences. The Yanomami dataset comprised 78 individuals from three villages along the Marauiá River (Amazonas, Brazil): Bicho-Açu (n = 31), the village closest to urban centers; Curuá (n = 20), with intermediate geographic remoteness and urban influence; and Xamakorona (n = 27), the most remote population examined (**Fig. 1a**). The urban cohort comprised 39 participants recruited in Belgium, with recruitment aimed at obtaining age and sex representation broadly comparable to that of the Yanomami participants. Population-level urbanization scores were calculated as described by McCall et al. (2019)^23^. Demographic characteristics, geographic reference coordinates, and urbanization scores are provided in **Supplementary Table 1**. Fecal samples were collected in sterile sampling bags for Yanomami participants or in Fecotainer® containers for Belgian participants and were immediately aliquoted into sterile 50-ml tubes. During fieldwork, Yanomami samples were kept at approximately 4°C and maintained under anaerobic conditions using the BD GasPak™ EZ Anaerobe Pouch System (Becton, Dickinson and Company), then transported to the laboratory, and stored frozen until processing.

### DNA extraction and sequencing

DNA extraction and shotgun metagenomic sequencing were performed by CosmosID (USA). Genomic DNA was extracted from fecal material using the DNeasy PowerSoil Pro Kit (QIAGEN) according to the manufacturer’s instructions. DNA concentration was quantified fluorometrically using the Qubit system (Thermo Fisher Scientific). Libraries were sequenced as paired-end 2×150 bp reads on the Element AVITI and Illumina NovaSeq X Plus platforms.

### Data preprocessing

Raw read quality was assessed with FastQC v0.12.1^54^ and summarized across samples with MultiQC v1.29^55^. Adapter trimming was performed in two sequential rounds with Atropos v1.1.32^56^ to remove Illumina universal adapter sequence (AGATCGGAAGAG) and Nextera transposase sequence (CTGTCTCTTATA). Additional quality filtering was performed with fastp v0.23.2^57^ to remove low-complexity reads (--complexity_threshold 30), trim poly-G tails (--trim_poly_g), remove the first 10 bases from both mates (--trim_front1 10, --trim_front2 10), and trim low-quality bases from the 3′ end using a sliding-window approach (--cut_tail, --cut_tail_window_size 3, --cut_tail_mean_quality 25). Reads shorter than 90 bp after trimming or with mean quality < Q25 were discarded. Host-derived reads were removed by aligning quality-filtered reads against the human reference genome GRCh38 using Bowtie2 v2.5.4^58^ in --very-sensitive-local mode; only unmapped read pairs were retained for downstream analysis.

### Metagenome assembly

High-quality, host-depleted reads were co-assembled separately for each population using metaSPAdes v3.15.5^59^ in metagenomic mode (--meta). Population-specific co-assemblies were used to improve recovery of the shared gene repertoire within each population while limiting cross-population assembly complexity^60^. Assembly quality was evaluated by remapping preprocessed reads to the corresponding co-assembly with BWA-MEM v0.7.17^61^ and summarizing assembly statistics with MetaQUAST v5.3.0^62^. Abundance was then estimated with kallisto v0.46.1^63^ in quant mode against the contigs catalog, generating raw counts and counts per million (CPM). CPM-length-normalized values were used for descriptive analyses, whereas raw counts were retained for differential abundance testing.

### Taxonomic assignment, gene prediction, and functional annotation

Assembled contigs were taxonomically assigned using MMseqs2 taxonomy v16.747c6^64^ using the Genome Taxonomy Database (GTDB; v226, released Apr 16, 2025)^65^ to assign bacterial and archaeal sequences. Contigs not assigned to Bacteria or Archaea were subsequently queried against the NCBI non-redundant protein database (nr; download date: Jul 18, 2025) to recover broader taxonomic assignments from other domains of life. Full lineage output was generated with --tax-lineage 1, and taxonomic strings were reformatted with TaxonKit^66^ using reformat2.

Protein-coding genes were predicted from each co-assembly with Prokka v1.14.6^67^ using the --metagenome option, which is appropriate for fragmented metagenomic assemblies. Functional annotation of predicted coding sequences was performed with eggNOG-mapper v2.1.12^68^ against eggNOG database v5.0.2^69^ in DIAMOND mode using --sensmode ultra-sensitive and an e-value cutoff of 1e-5. KEGG Orthologs (KOs) assignments and their corresponding KEGG hierarchical annotations were retained for downstream functional profiling^70^. Carbohydrate-active enzymes (CAZymes) were annotated using run_dbcan from dbCAN3^71^, combining HMMER, dbCAN_sub, and DIAMOND searches with default coverage thresholds and an e-value cutoff of 1e-5. CAZyme annotations were retained only when supported by at least two of the three dbCAN tools. To infer preferred carbohydrate substrates, retained CAZymes were additionally compared against the CAZyme-substrate annotations available in the Cayman database v0.10.1^47^.

### Feature filtering

All count matrices and associated annotations were imported into R v4.3.2^72^ for downstream analyses. To reduce sparsity and minimize the influence of likely spurious low-abundance features, filtering was performed independently for each annotation type: taxonomy, KEGG orthologs, and CAZymes. For each dataset, counts were collapsed to the smallest non-redundant annotation unit, namely TaxID for taxonomic annotations, eggNOG ortholog ID for KEGG-based analyses, and gene ID for CAZyme annotations. Features were retained only when they reached at least 1 CPM in at least 10% of samples from their corresponding population.

Taxonomic analyses were managed with phyloseq v1.46.0^73^, with additional filtering performed using metagMisc v0.5.0^74^. Taxonomic summaries from phylum to genus were generated from genus-resolved data and then aggregated to the desired rank. For composition plots, genera were first subjected to sample-wise abundance trimming with a minimum relative abundance of 0.1%, using phyloseq_filter_sample_wise_abund_trim. A second prevalence filter was then applied with phyloseq_filter_prevalence, retaining genera present in >10% of samples. For taxonomic feature-centered taxonomic analyses, including alpha and beta diversity, an abundance threshold of 0.01% was used, followed by the same prevalence criterion.

### Taxonomic profiling and ecological diversity

Taxonomic composition was summarized as relative abundance at the phylum and genus levels. Taxa shared among populations were visualized using Venn diagrams constructed with the ggvenn v0.1.16^75^. A prevalence threshold of ≥ 50% of samples per population was applied to define taxon presence within each group. Alpha diversity was calculated from filtered taxonomic tables using observed richness and Shannon diversity, with the alpha function in the microbiome package v1.24.0^76^. Beta diversity was assessed using Bray-Curtis dissimilarity matrices computed in phyloseq, and principal coordinate analysis (PCoA) was used for ordination.

### Co-occurrence network analysis

Genus-level co-occurrence networks were inferred separately for each population from filtered abundance tables. Pairwise Pearson correlations were calculated with corr.test in psych v2.5.6^77^, and only statistically significant associations with |*r*| ≥ 0.5 and FDR-adjusted *P* ≤ 0.05 were retained as edges. Network topology was characterized using igraph v2.1.4^78^, including node degree and centrality measures. Modules were detected with the Louvain community detection algorithm^79^. Node roles were classified according to within-module connectivity (Z_i_) and between-module connectivity (P_i_) as modular hubs (Z_i_ > 2.5, P_i_ < 0.62), connectors (Z_i_ < 2.5, P_i_ > 0.62), or peripherals (Z_i_ ≤ 2.5, P_i_ ≤ 0.62). Candidate hub taxa were additionally identified from the five highest Kleinberg hub scores^80^. Network robustness was evaluated by targeted attack simulations in NetSwan v0.1^81^. In this method, nodes were removed iteratively in descending order of degree, and the resulting loss of connectivity was tracked.

### Functional Profiling

Functional profiling was focused on genes assigned to KEGG orthologs to characterize differences in microbial metabolic potential across populations. KO profiles were aggregated at the KEGG metabolism subclass level for broad functional comparisons and at the metabolic pathway level for biomarker discovery and targeted analyses. Shared features, alpha- and beta-diversity analyses based on metabolic KO profiles followed the same framework described for taxonomic profiles. In parallel, carbohydrate-utilization potential was characterized using CAZyme annotations and their predicted preferred substrates, with CAZyme- and substrate-based profiles compared across populations using the same diversity framework.

To further characterize microbial butyrate biosynthetic potential, we reconstructed the four major butyrate production pathways described for the human gut microbiome: the acetyl-CoA, succinate, glutamate, and lysine pathways^45,82^. KOs were selected based on their association with the enzyme commission (EC) numbers corresponding to the enzymatic reactions comprising these pathways.

### Statistical Analyses

All statistical analyses were performed in R. Overall differences among populations were assessed using the Kruskal-Wallis test. When significant, pairwise comparisons were performed using two-sided Wilcoxon rank-sum tests. *P* values were adjusted for multiple comparisons using the Benjamini-Hochberg procedure to control the false discovery rate (FDR)^83^, with FDR-adjusted *P* < 0.05 considered statistically significant and values between 0.05 and 0.1 considered trends. Multivariate differences in taxonomic and functional composition were assessed by PERMANOVA using adonis2 in vegan v2.7-2^84^ on Bray-Curtis dissimilarity matrices with 999 permutations. Post-hoc pairwise PERMANOVA comparisons were performed using the same number of permutations and significance threshold.

Differential abundance of genera was assessed using MaAsLin2 v1.16.0^85^. Genus relative-abundance profiles were total-sum scaled (TSS), and population was included as a fixed effect, and age group and sex were included as random effects in all contrasts. Associations were considered significant when the absolute regression coefficient (|β|) was >1 and the Benjamini-Hochberg-adjusted *q* value was < 0.05. Differential abundance of individual KOs and CAZymes was assessed using DESeq2 v1.34.0^86^ on rounded raw count matrices. Pairwise models were used for the population contrasts of interest, and features were considered differentially abundant when they met both an absolute log2 fold-change > 1 and an FDR-adjusted *P* value < 0.05.

Biomarker analysis of KOs aggregated at the KEGG Metabolism pathway level was performed using LEfSe^87^, implemented through microbiomeMarker v1.13.2^88^. Statistical significance was determined after Benjamini-Hochberg adjustment of LEfSe *P* values, and pathways were retained as biomarkers when they showed an LDA effect size > 2 and an adjusted *P* value < 0.05.

Over-representation analysis (ORA) was used to determine whether functional or substrate categories were enriched among differentially abundant features. Analyses were performed separately for features enriched in each direction of a given population contrast. For each comparison, the background universe consisted of all features included in the corresponding differential-abundance analysis that could be assigned to the annotation level under evaluation, whereas the query set comprised the differentially abundant features enriched in the population of interest. Feature-to-category relationships were supplied as custom annotation sets to the enricher function in clusterProfiler v4.21.1^89^. Enrichment was evaluated using a hypergeometric test. *P* values were adjusted for multiple testing using the Benjamini–Hochberg FDR procedure, and categories with adjusted *P* ≤ 0.05 were considered significantly overrepresented.

## Data availability

Individual-level metagenomic data generated in this study are available upon request, respecting Indigenous data sovereignty and ownership and complying with applicable Brazilian regulations governing genetic resources and benefit sharing. Requests should outline the proposed uses and scientific objectives, explain how data security and participant privacy will be protected, and describe the potential benefits for the Yanomami community.

## Acknowledgements

We thank all study volunteers, the Yanomami communities and leaders for their contribution to this research. We thank the “Secretaria de Saúde Indígena” (SESAI), “Fundação Nacional dos Povos Indígenas” (FUNAI) and “Serviço e Cooperação com Povo Yanomami” (SECOYA) for supporting engagement with the participating communities. We thank Dirk Van Dijck for logistical support during fieldwork. We thank COPAN for sponsoring the sampling material. We acknowledge the Multiuser Computational Center of the Federal University of ABC (CCM-UFABC) for providing the high-performance computing resources used in this work.

## Contributions

L.A.L.C.: data curation, formal analysis, investigation, methodology, software, visualization, writing – original draft, writing – review & editing. M.T.: formal analysis, investigation, methodology, writing – review & editing. M.C.: formal analysis, writing – review & editing. C.C., D.G. and E.A.-V.: methodology, writing – review & editing. L.V. and L.C.P.: conceptualization, funding acquisition, investigation, project administration, resources, supervision, writing – review & editing. All authors reviewed and approved the final manuscript.

