## Supplementary Material for "Comparisons among Yanomami populations reveal transitional gut microbiome configurations under urban influence"

**Supplementary Figure 1. Extended taxonomic characterization and evaluation of demographic covariates. a,** Percentage of sequences assigned at phylum, genus, and species levels. Mean ± SD classification rates at the phylum level were BEL, 96.72 ± 0.30%; BIC, 96.89 ± 0.30%; CUR, 96.89 ± 0.44%; XAM, 97.17 ± 0.27%; at genus level, 81.28 ± 1.95%, 83.67 ± 1.34%, 84.94 ± 2.19%, and 85.09 ± 1.53%; and at species level, 56.25 ± 3.29%, 56.95 ± 4.54%, 55.40 ± 5.50%, and 53.57 ± 5.01%, respectively. **b,** Mean ± SD relative abundance of taxonomic domains before abundance/prevalence filtering. Archaea: BEL, 0.125 ± 0.243%; BIC, 0.268 ± 0.310%; CUR, 0.291 ± 0.513%; XAM, 0.304 ± 0.524% (Kruskal–Wallis $H$ = 10.71, FDR-adjusted $P$ = 0.013). Bacteria: 99.719 ± 0.278%, 99.092 ± 0.825%, 99.427 ± 0.609%, and 99.473 ± 0.749%, respectively ($H$ = 21.50, $P$ < 0.001). Eukaryota: 0.017 ± 0.030%, 0.418 ± 0.543%, 0.071 ± 0.144%, and 0.110 ± 0.201% ($H$ = 37.02, $P$ < 0.001). Viruses: 0.139 ± 0.081%, 0.222 ± 0.085%, 0.211 ± 0.081%, and 0.114 ± 0.057% ($H$ = 33.71, $P$ < 0.001). **c,** Per-sample phylum-level relative-abundance profiles, with the five most abundant phyla shown individually and remaining taxa grouped as in **Figure 1**; samples are ordered by Bacillota abundance. **d,** Per-sample genus-level relative-abundance profiles using the representation scheme described in **Figure 1**; samples are ordered by *Prevotella* abundance. **e,f,** PCoA of Bray–Curtis taxonomic dissimilarities evaluated in relation to potential demographic covariates. Density plots show the distribution of sample scores along PCoA1 (top) and PCoA2 (right), grouped according to the color coding in each panel. No population-by-sex interaction was detected (**e**; PERMANOVA: $R^2$ = 0.018, $F$ = 0.943, $P$ = 0.519), nor a population-by-age-group interaction (**f**; $R^2$ = 0.062, $F$ = 0.869, $P$ = 0.840). Significance for FDR-adjusted pairwise comparisons, where shown, is ***$P$ < 0.001, **$P$ < 0.01, *$P$ < 0.05, and ~$P$ < 0.1.

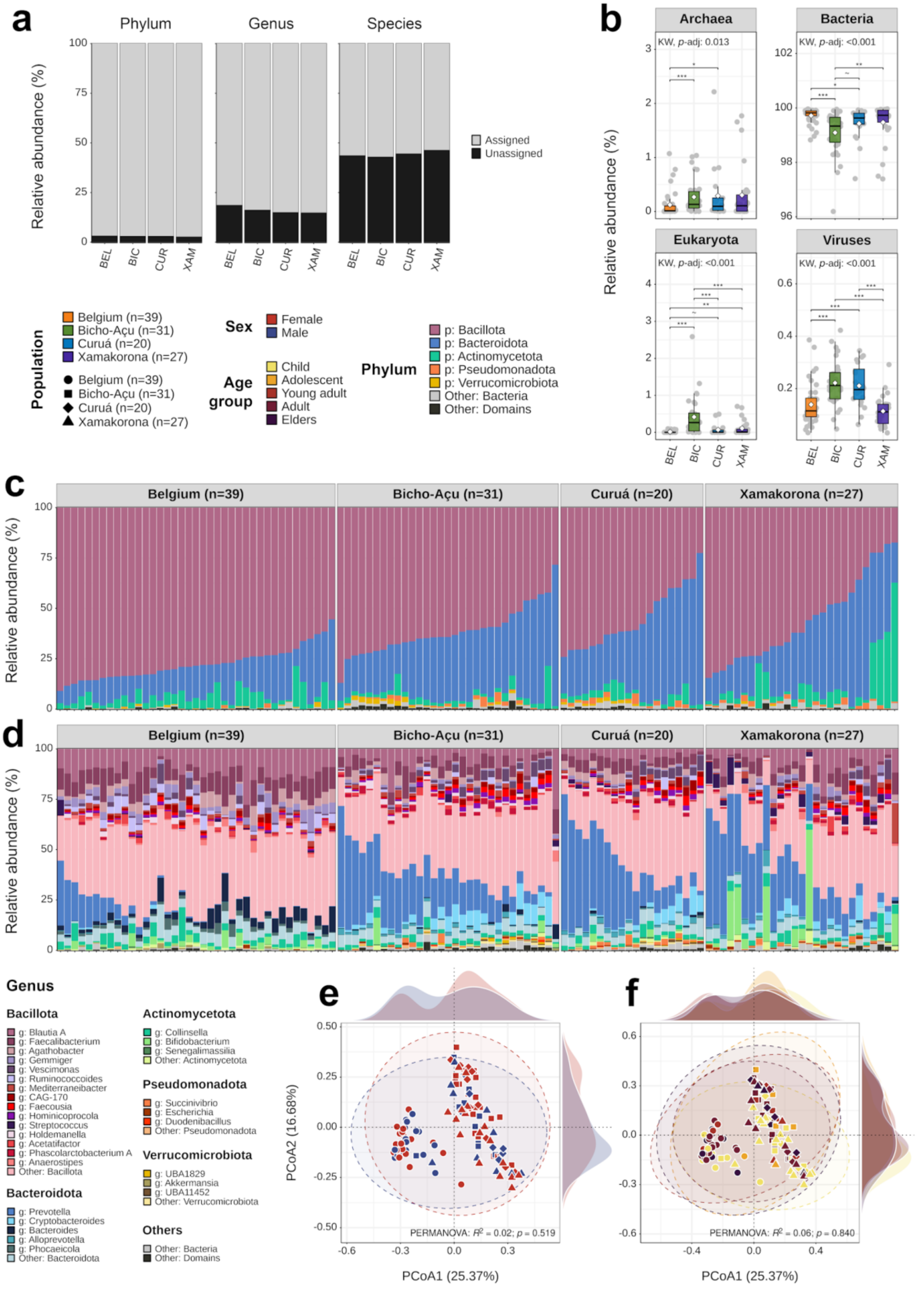

a
Phylum
Genus
Species
Relative abundance (%)
Assigned
Unassigned
BEL
BIC
CUR
XAM
b
Archaea
Bacteria
Eukaryota
Viruses
KW, p-adj: 0.013
KW, p-adj: <0.001
Relative abundance (%)
Population
Belgium (n=39)
Bicho-Açu (n=31)
Curuá (n=20)
Xamakorona (n=27)
Sex
Female
Male
Age group
Child
Adolescent
Young adult
Adult
Elders
Phylum
p: Bacillota
p: Bacteroidota
p: Actinomycetota
p: Pseudomonadota
p: Verrucomicrobiota
Other: Bacteria
Other: Domains
c
d
Genus
Bacillota
g: Blautia A
g: Faecalibacterium
g: Agathobacter
g: Gemmiger
g: Vescimonas
g: Ruminococcoides
g: Mediterraneibacter
g: CAG-170
g: Faecousia
g: Hominicoprocola
g: Streptococcus
g: Holdemanella
g: Acetatifactor
g: Phascolarctobacterium A
g: Anaerostipes
Other: Bacillota
Bacteroidota
g: Prevotella
g: Cryptobacteroides
g: Bacteroides
g: Alloprevotella
g: Phocaeicola
Other: Bacteroidota
Actinomycetota
g: Collinsella
g: Bifidobacterium
g: Senegalimassilia
Other: Actinomycetota
Pseudomonadota
g: Succinivibrio
g: Escherichia
g: Duodenibacillus
Other: Pseudomonadota
Verrucomicrobiota
g: UBA1829
g: Akkermansia
g: UBA11452
Other: Verrucomicrobiota
Others
Other: Bacteria
Other: Domains
e
f
PCoA2 (16.68%)
PCoA1 (25.37%)
PERMANOVA: R² = 0.02; p = 0.519
PERMANOVA: R² = 0.06; p = 0.840

**Supplementary Figure 2. Metabolic functional differences among Yanomami populations. a,** Assignment of differentially abundant KOs identified by DESeq2 in BIC–CUR, BIC–XAM, and CUR–XAM comparisons to functional subclasses within the KEGG Metabolism class. Colored bars represent differentially abundant KOs and gray bars non-differentially abundant KOs. Triangles indicate subclasses significantly enriched among differentially abundant KOs by over-representation analysis using a hypergeometric test with Benjamini–Hochberg-adjusted $P \leq 0.05$. Individual KOs were considered differentially abundant at $|\log_2$ fold-change$| > 1$ and FDR-adjusted $P < 0.05$. **b,** KEGG metabolic pathways evaluated by LEfSe for the same Yanomami comparisons. Pathways were retained as significant biomarkers at LDA effect size > 2 and Benjamini–Hochberg-adjusted $P < 0.05$. The annotation bar adjacent to pathway names indicates the metabolism subclass defined in **a**. Significance, where indicated, follows ***FDR-adjusted $P < 0.001$, **$P < 0.01$, *$P < 0.05$, and ~$P < 0.1$.

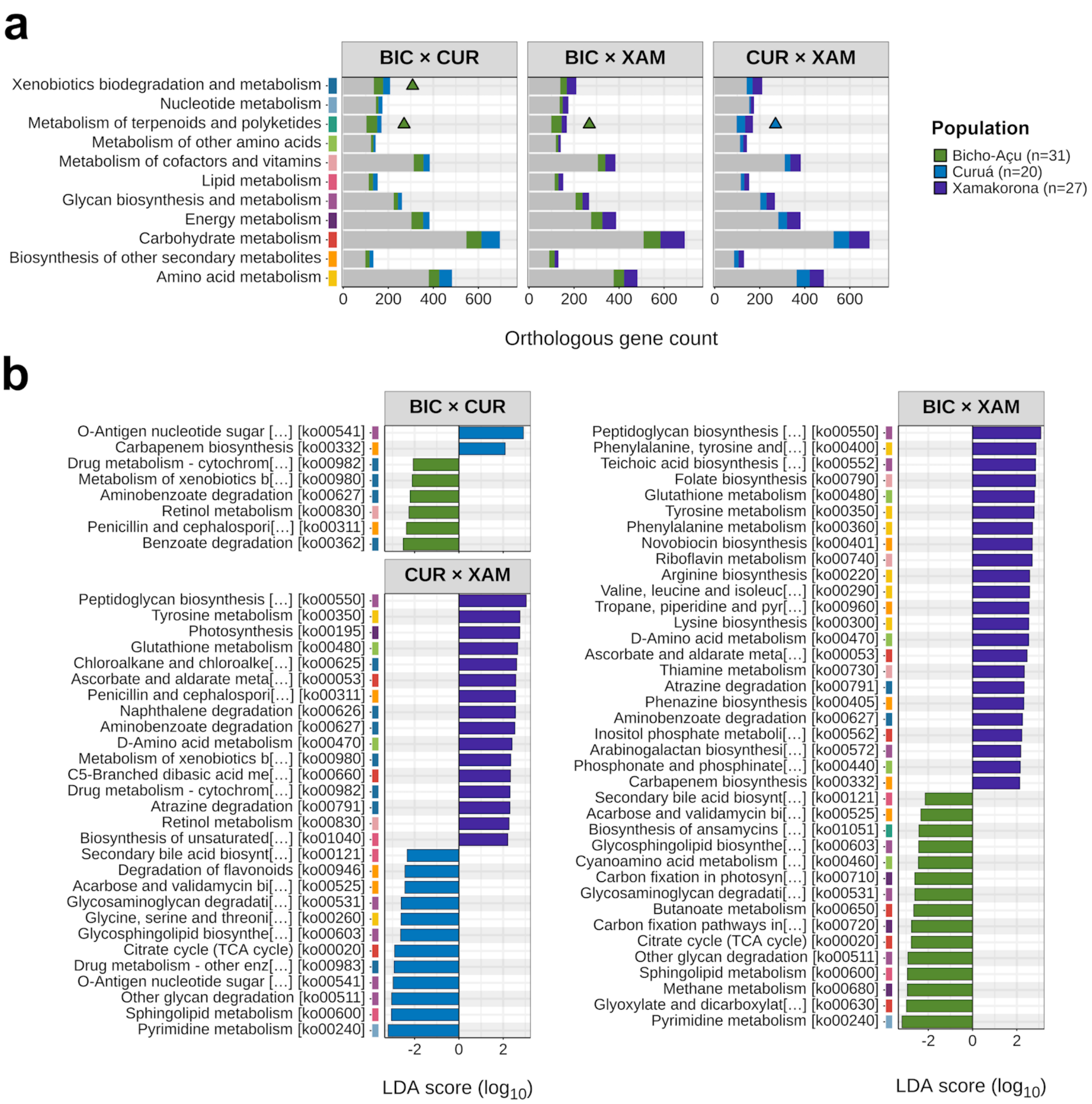
a
BIC × CUR
BIC × XAM
CUR × XAM
Xenobiotics biodegradation and metabolism
Nucleotide metabolism
Metabolism of terpenoids and polyketides
Metabolism of other amino acids
Metabolism of cofactors and vitamins
Lipid metabolism
Glycan biosynthesis and metabolism
Energy metabolism
Carbohydrate metabolism
Biosynthesis of other secondary metabolites
Amino acid metabolism
0 200 400 600
Orthologous gene count
Population
Bicho-Açu (n=31)
Curuá (n=20)
Xamakorona (n=27)
b
BIC × CUR
O-Antigen nucleotide sugar [...] [ko00541]
Carbapenem biosynthesis [ko00332]
Drug metabolism - cytochrom[...] [ko00982]
Metabolism of xenobiotics b[...] [ko00980]
Aminobenzoate degradation [ko00627]
Retinol metabolism [ko00830]
Penicillin and cephalospori[...] [ko00311]
Benzoate degradation [ko00362]
CUR × XAM
Peptidoglycan biosynthesis [...] [ko00550]
Tyrosine metabolism [ko00350]
Photosynthesis [ko00195]
Glutathione metabolism [ko00480]
Chloroalkane and chloroalke[...] [ko00625]
Ascorbate and aldarate meta[...] [ko00053]
Penicillin and cephalospori[...] [ko00311]
Naphthalene degradation [ko00626]
Aminobenzoate degradation [ko00627]
D-Amino acid metabolism [ko00470]
Metabolism of xenobiotics b[...] [ko00980]
C5-Branched dibasic acid me[...] [ko00660]
Drug metabolism - cytochrom[...] [ko00982]
Atrazine degradation [ko00791]
Retinol metabolism [ko00830]
Biosynthesis of unsaturated[...] [ko01040]
Secondary bile acid biosynt[...] [ko00121]
Degradation of flavonoids [ko00946]
Acarbose and validamycin bi[...] [ko00525]
Glycosaminoglycan degradati[...] [ko00531]
Glycine, serine and threoni[...] [ko00260]
Glycosphingolipid biosynthe[...] [ko00603]
Citrate cycle (TCA cycle) [ko00020]
Drug metabolism - other enz[...] [ko00983]
O-Antigen nucleotide sugar [...] [ko00541]
Other glycan degradation [ko00511]
Sphingolipid metabolism [ko00600]
Pyrimidine metabolism [ko00240]
-2 0 2
LDA score (log10)
BIC × XAM
Peptidoglycan biosynthesis [...] [ko00550]
Phenylalanine, tyrosine and[...] [ko00400]
Teichoic acid biosynthesis [...] [ko00552]
Folate biosynthesis [ko00790]
Glutathione metabolism [ko00480]
Tyrosine metabolism [ko00350]
Phenylalanine metabolism [ko00360]
Novobiocin biosynthesis [ko00401]
Riboflavin metabolism [ko00740]
Arginine biosynthesis [ko00220]
Valine, leucine and isoleuc[...] [ko00290]
Tropane, piperidine and pyr[...] [ko00960]
Lysine biosynthesis [ko00300]
D-Amino acid metabolism [ko00470]
Ascorbate and aldarate meta[...] [ko00053]
Thiamine metabolism [ko00730]
Atrazine degradation [ko00791]
Phenazine biosynthesis [ko00405]
Aminobenzoate degradation [ko00627]
Inositol phosphate metaboli[...] [ko00562]
Arabinogalactan biosynthesi[...] [ko00572]
Phosphonate and phosphinate[...] [ko00440]
Carbapenem biosynthesis [ko00332]
Secondary bile acid biosynt[...] [ko00121]
Acarbose and validamycin bi[...] [ko00525]
Biosynthesis of ansamycins [...] [ko01051]
Glycosphingolipid biosynthe[...] [ko00603]
Cyanoamino acid metabolism [...] [ko00460]
Carbon fixation in photosyn[...] [ko00710]
Glycosaminoglycan degradati[...] [ko00531]
Butanoate metabolism [ko00650]
Carbon fixation pathways in[...] [ko00720]
Citrate cycle (TCA cycle) [ko00020]
Other glycan degradation [ko00511]
Sphingolipid metabolism [ko00600]
Methane metabolism [ko00680]
Glyoxylate and dicarboxylat[...] [ko00630]
Pyrimidine metabolism [ko00240]
-2 0 2
LDA score (log10)

**Supplementary Figure 3. Population-specific differential abundance of KOs contributing to microbial butyrate biosynthesis.** Pairwise DESeq2 comparisons are shown for the KOs highlighted in the butyrate pathways in **Figure 6**. Differential abundance was defined as Benjamini–Hochberg FDR-adjusted $P < 0.05$. Statistical significance for each pairwise contrast is indicated as ***$P < 0.001$, **$P < 0.01$, *$P < 0.05$, and ~$P < 0.1$.

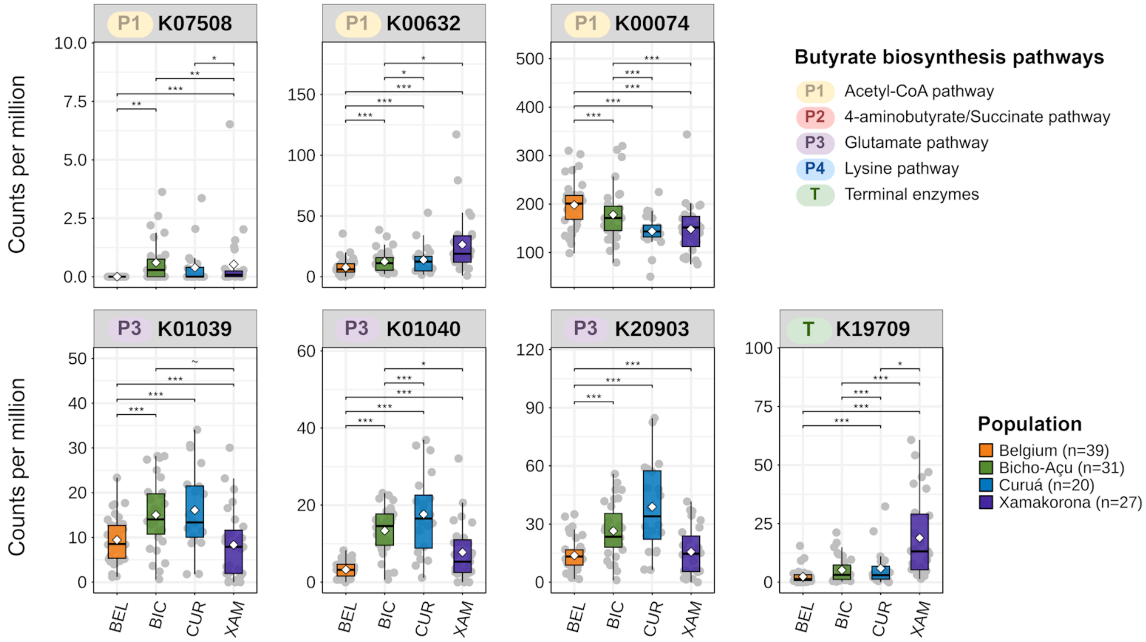

**Supplementary Figure 4. Predicted carbohydrate-substrate profiles and their taxonomic contributors differ across populations. a,** Mean genus-level taxonomic contribution to CAZymes assigned to Cayman level-2 substrate categories, including non-starch polysaccharides (NSP), resistant oligosaccharides (RO), resistant starch (RS), mucin (MUC), and peptidoglycan (PG). The 10 most abundant genus-level contributors are shown for each population and substrate category. **b,** Relative abundance of CAZymes assigned to the 10 most abundant individual Cayman substrates. Mean ± SD values for BEL, BIC, CUR, and XAM, respectively, were: xylan, 7.540 ± 0.376%, 6.601 ± 0.531%, 6.694 ± 0.681%, and 6.797 ± 0.486% (Kruskal–Wallis $H$ = 50.41, $P < 0.001$); mucin glycan, 6.193 ± 0.577%, 6.919 ± 0.703%, 6.625 ± 0.617%, and 6.379 ± 0.595% ($H$ = 25.57, $P < 0.001$); arabinoxylan, 6.446 ± 0.449%, 5.418 ± 0.526%, 5.659 ± 0.726%, and 5.756 ± 0.590% ($H$ = 46.27, $P < 0.001$); xyloglucan, 5.870 ± 0.241%, 5.635 ± 0.296%, 5.887 ± 0.429%, and 5.555 ± 0.223% ($H$ = 25.40, $P < 0.001$); starch, 5.170 ± 0.660%, 5.803 ± 0.700%, 5.499 ± 0.872%, and 6.238 ± 0.958% ($H$ = 25.45, $P < 0.001$); galactan, 5.437 ± 0.251%, 5.593 ± 0.200%, 5.807 ± 0.384%, and 5.755 ± 0.275% ($H$ = 25.75, $P < 0.001$); glucomannan, 5.628 ± 0.300%, 5.344 ± 0.281%, 5.606 ± 0.356%, and 5.362 ± 0.240% ($H$ = 19.63, $P < 0.001$); β-glucan, 5.585 ± 0.284%, 5.402 ± 0.338%, 5.583 ± 0.420%, and 5.305 ± 0.251% ($H$ = 15.13, $P = 0.002$); arabinan, 5.757 ± 0.322%, 4.750 ± 0.396%, 4.845 ± 0.445%, and 4.685 ± 0.670% ($H$ = 60.73, $P < 0.001$); arabinogalactan, 5.718 ± 0.357%, 4.701 ± 0.428%, 4.759 ± 0.356%, and 4.471 ± 0.515% ($H$ = 70.97, $P < 0.001$). Significant omnibus tests were followed by pairwise two-sided Wilcoxon rank-sum tests with Benjamini–Hochberg FDR correction. Significance for FDR-adjusted pairwise comparisons is indicated as ***$P < 0.001$, **$P < 0.01$, *$P < 0.05$, and ~$P < 0.1$.

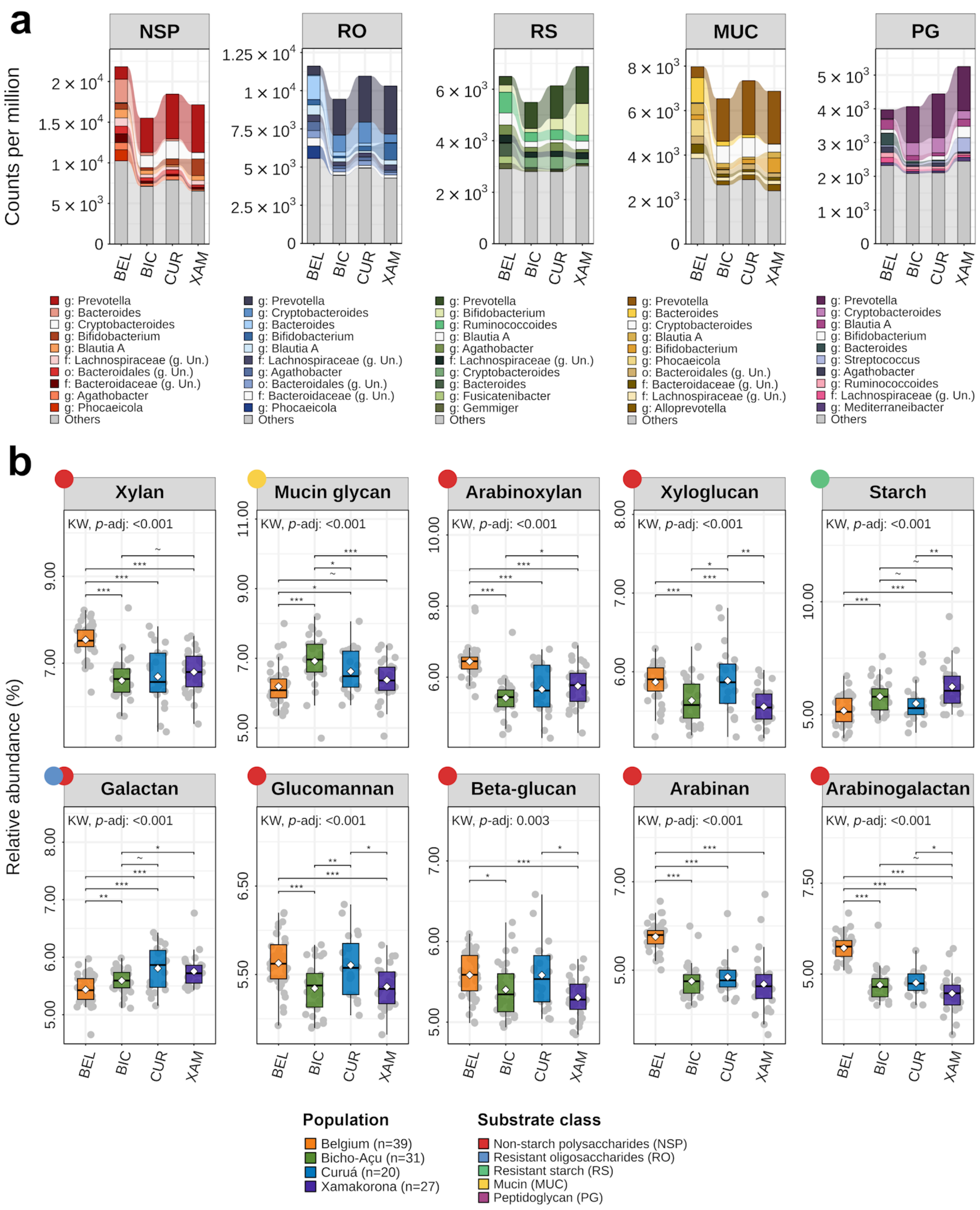
a
NSP
RO
RS
MUC
PG
Counts per million
BEL
BIC
CUR
XAM
g: Prevotella
g: Bacteroides
g: Cryptobacteroides
g: Bifidobacterium
g: Blautia A
f: Lachnospiraceae (g. Un.)
o: Bacteroidales (g. Un.)
f: Bacteroidaceae (g. Un.)
g: Agathobacter
g: Phocaeicola
Others
g: Prevotella
g: Cryptobacteroides
g: Bacteroides
g: Bifidobacterium
g: Blautia A
f: Lachnospiraceae (g. Un.)
g: Agathobacter
o: Bacteroidales (g. Un.)
f: Bacteroidaceae (g. Un.)
g: Phocaeicola
Others
g: Prevotella
g: Bifidobacterium
g: Ruminococcoides
g: Blautia A
g: Agathobacter
f: Lachnospiraceae (g. Un.)
g: Cryptobacteroides
g: Bacteroides
g: Fusicatenibacter
g: Gemmiger
Others
g: Prevotella
g: Bacteroides
g: Cryptobacteroides
g: Blautia A
g: Bifidobacterium
g: Phocaeicola
o: Bacteroidales (g. Un.)
f: Bacteroidaceae (g. Un.)
f: Lachnospiraceae (g. Un.)
g: Alloprevotella
Others
g: Prevotella
g: Cryptobacteroides
g: Blautia A
g: Bifidobacterium
g: Bacteroides
g: Streptococcus
g: Agathobacter
g: Ruminococcoides
f: Lachnospiraceae (g. Un.)
g: Mediterraneibacter
Others
b
Relative abundance (%)
Xylan
Mucin glycan
Arabinoxylan
Xyloglucan
Starch
Galactan
Glucomannan
Beta-glucan
Arabinan
Arabinogalactan
KW, p-adj: <0.001
KW, p-adj: 0.003
Population
Belgium (n=39)
Bicho-Açu (n=31)
Curuá (n=20)
Xamakorona (n=27)
Substrate class
Non-starch polysaccharides (NSP)
Resistant oligosaccharides (RO)
Resistant starch (RS)
Mucin (MUC)
Peptidoglycan (PG)

**Supplementary Table 1. Characteristics of the study populations.** Demographic characteristics and geographic reference coordinates of the Belgian urban cohort (n = 39) and Yanomami subpopulations Bicho-Açu (n = 31), Curuá (n = 20), and Xamakorona (n = 27).

| | **Belgium (n = 39)** | **Bicho-Açu (n = 31)** | **Curuá (n = 20)** | **Xamakorona (n = 27)** |
|---|---|---|---|---|
| **Sex**, n (%) | | | | |
| Female | 21 (53.85) | 21 (67.74) | 13 (65) | 17 (62.96) |
| Male | 18 (46.15) | 10 (32.26) | 7 (35) | 10 (37.04) |
| **Age group**, n (%) | | | | |
| Child (2-12y) | 10 (25.64) | 8 (25.81) | 7 (35) | 14 (51.85) |
| Adolescent (13-17y) | 3 (7.69) | 6 (19.35) | - | 1 (3.7) |
| Young adult (18-25y) | 6 (15.38) | 4 (12.9) | 2 (10) | 2 (7.41) |
| Adult (26-50y) | 14 (35.9) | 12 (38.71) | 7 (35) | 7 (25.93) |
| Elder (>51y) | 6 (15.38) | 1 (3.23) | 4 (20) | 3 (11.11) |
| **Localities** | | | | |
| Urbanization score[1] | > 90 | 54.75 | 36 | 2.5 |
| Coordinates[2] | 51°01'28.4"N 3°43'30.8"E | 0°18'53.760"S 65°07'07.180"W | 0°12'47.381"S 65°04'39.950"W | 1°00'55.459"N 65°03'12.010"W |

[1]Urbanization score was used to quantitatively characterize the degree of lifestyle transition across the Yanomami villages, following an established scoring framework adapted from McCall et al. (2019). This index incorporates access to industrialized foods, use of personal hygiene products, access to healthcare services, and previous exposure to antibiotics. [2]Coordinates (latitude/longitude) correspond to reference locations for the participant community/city. They represent community-level reference points rather than individual household locations.

McCall, Laura-Isobel, et al. "Home chemical and microbial transitions across urbanization". *Nature microbiology* 5.1 (2019): 108-115. https://doi.org/10.1038/s41564-019-0593-4

**Supplementary Table 2. Metagenomic sequencing, preprocessing, assembly, and read-mapping statistics.** Sequencing and preprocessing statistics include mean raw reads per sample and reads retained after quality filtering, adapter removal, and human-read depletion. Population-level co-assembly statistics include contig number (≥500 bp), total assembled length for contigs ≥500 bp, largest contig, N50, and L50. Mapping statistics report the percentage of reads mapped to the corresponding population co-assembly overall and at high mapping quality (HQ; mapping quality ≥Q20). Summary values are reported as mean ± SD where applicable.

| **Preprocessing statistics** | **Belgium (n = 39)** | **Bicho-Açu (n = 31)** | **Curuá (n = 20)** | **Xamakorona (n = 27)** |
|---|---|---|---|---|
| Raw read count | 8,502,520.36 (±2,344,198.25) | 7,819,622.58 (±2,052,053.02) | 7,276,987.75 (±1,680,090.39) | 11,265,194.88 (±3,630,740.09) |
| Retained after QC | 8,187,045.46 (±2,229,140.63) | 6,707,782.84 (±1,743,113.37) | 6,236,703.40 (±1,256,485.30) | 10,263,958.94 (±3,457,616.19) |
| **Assembly statistics** | | | | |
| # of contigs | 455,762 | 868,170 | 621,685 | 933,726 |
| Total length | 818,754,456 | 1,580,739,039 | 1,102,105,401 | 1,694,966,485 |
| Largest contig | 353,470 | 228,903 | 303,954 | 345,887 |
| N50 | 2,767 | 2,793 | 2,671 | 2,782 |
| L50 | 59,620 | 118,112 | 83,378 | 126,846 |
| **Mapping statistics** | | | | |
| Mapping rate (any, %) | 97.37 (±1.44) | 98.35 (±0.7) | 97.43 (±1.19) | 97.76 (±2.99) |
| Mapping rate (HQ, %) | 92.45 (±2.41) | 92.52 (±0.77) | 91.71 (±1.27) | 90.16 (±3.09) |

**Supplementary Table 3. Gene prediction and functional annotation across databases.** Summary of predicted coding sequences generated using Prokka; genes assigned to eggNOG orthologous groups and retained after abundance/prevalence filtering; genes assigned to KEGG orthologs and KEGG Metabolism subclasses; and carbohydrate-active enzyme (CAZymes) annotations.

| **Feature** | **Belgium (n = 39)** | **Bicho-Açu (n = 31)** | **Curuá (n = 20)** | **Xamakorona (n = 27)** |
|---|---|---|---|---|
| (i) Predicted CDS (Coding Sequences) | 784,200 | 1,518,897 | 1,058,855 | 1,622,780 |
| (ii) Annotated[1] and filtered[2] genes | 511,729 | 896,123 | 691,557 | 869,474 |
| (iii) Genes annotated w/ KO | 318,030 | 585,465 | 454,521 | 565,823 |
| Genes annotated w/ KO metabolism | 115,471 | 211,418 | 166,609 | 206,481 |
| (iv) Genes annotated as CAZyme | 21,411 | 36,870 | 25,775 | 39,991 |

[1] Annotated with EggNOG mapper. [2] Genes with >1 CPM in at least 10% of samples within each group.

**Supplementary Table 4. PERMANOVA of taxonomic beta diversity based on Bray–Curtis dissimilarities.** PERMANOVA results testing differences in taxonomic composition among Belgium (n = 39), Bicho-Açu (n = 31), Curuá (n = 20), and Xamakorona (n = 27). Pairwise post-hoc PERMANOVA contrasts are reported with *F* statistics and Benjamini–Hochberg FDR-adjusted *P* values. All tests used 999 permutations.

| **Comparison** | **Source** | **df** | **SS** | **$R^2$** | ***F*** | ***P*-adj** |
|---|---|---|---|---|---|---|
| Population | Model | 3 | 5.41 | 0.27 | 13.62 | <0.001 |
| | Residual | 113 | 14.95 | 0.73 | | |
| | Total | 116 | 20.36 | 1.00 | | |
| **PERMANOVA post-hoc test** | | | | | **F** | ***P*-adj** |
| Belgium vs Bicho-Açu | | | | | 24.44 | <0.001 |
| Belgium vs Curuá | | | | | 22.32 | <0.001 |
| Belgium vs Xamakorona | | | | | 22.14 | <0.001 |
| Bicho-Açu vs Curuá | | | | | 2.62 | 0.012 |
| Bicho-Açu vs Xamakorona | | | | | 4.66 | <0.001 |
| Curuá vs Xamakorona | | | | | 3.75 | 0.002 |

Abbreviations: df, degrees of freedom; SS, sum of squares; $R^2$, proportion of variance explained; *F*, PERMANOVA pseudo-F statistic.

**Supplementary Table 5. Genus-level co-occurrence network topology, node roles, and centrality metrics.** Population-specific network properties, including global topology and node-level centrality measures. Node roles were defined from within-module connectivity ($Z_i$) and among-module connectivity ($P_i$), with modules detected using the Louvain algorithm.

| **Metric** | **Belgium (n = 39)** | **Bicho-Açu (n = 31)** | **Curuá (n = 20)** | **Xamakorona (n = 27)** |
|---|---|---|---|---|
| Number of nodes | 115 | 182 | 165 | 184 |
| Number of edges | 236 | 665 | 749 | 1423 |
| Positives edges | 228 (96.61%) | 652 (98.05%) | 712 (95.06%) | 1404 (98.66%) |
| Negative edges | 8 (3.39%) | 13 (1.95%) | 37 (4.94%) | 19 (1.34%) |
| Modularity coefficient | 0.224 | 0 | 0.042 | 0.001 |
| Average degree[1] (KW *p*-adj. < 0.001) | 4.1 (±2.89) | 7.31 (±5.99) | 9.08 (±7.07) | 15.47 (±12.81) |
| Hub taxa[2] | *Vescimonas, Alistipes, Faecousia, Akkermansia, SFEL01* | *Faecousia, SFEL01, Merdousia, SFTH01, Evtepia* | *Limiplasma, Vescimonas, Butyricimonas, Firm-10, CAG-238* | *Faecivicinus, Vescimonas, SFTH01, CAG-238, Apopatocola* |
| Number of modules[3] | 15 | 10 | 11 | 10 |
| Modular hubs[4] | - | - | - | *g: Lachnospira* |
| Connectors[5] | 6 | 7 | 8 | 36 |
| Peripherals[6] | 109 | 175 | 157 | 147 |

[1] Means compared via Kruskal–Wallis/Wilcoxon rank-sum tests (FDR-adjusted *P*-value ≤ 0.05); [2] Hub taxa identified via Kleinberg's hubbiness score (top 5 shown); [3] Modules predicted using the Louvain method; [4] Modular hubs: $Z_i > 2.5$ and $P_i < 0.62$; [5] Connectors: $Z_i < 2.5$ and $P_i > 0.62$; [6] Peripherals: $Z_i \leq 2.5$ and $P_i \leq 0.62$.

**Supplementary Table 6. PERMANOVA of KEGG orthologs beta diversity based on Bray–Curtis dissimilarities.** PERMANOVA results testing differences in KEGG ortholog composition among Belgium (n = 39), Bicho-Açu (n = 31), Curuá (n = 20), and Xamakorona (n = 27). Pairwise post-hoc PERMANOVA contrasts are reported with *F* statistics and Benjamini–Hochberg FDR-adjusted *P* values. All tests used 999 permutations.

| **Comparison** | **Source** | **df** | **SS** | **$R^2$** | ***F*** | ***P*-adj** |
|---|---|---|---|---|---|---|
| Population | Model | 3 | 0.46 | 0.24 | 12.09 | <0.001 |
| | Residual | 113 | 1.43 | 0.76 | | |
| | Total | 116 | 1.89 | 1.00 | | |
| **PERMANOVA post-hoc test** | | | | | **F** | ***P*-adj** |
| Belgium vs Bicho-Açu | | | | | 20.89 | <0.001 |
| Belgium vs Curuá | | | | | 24.80 | <0.001 |
| Belgium vs Xamakorona | | | | | 17.33 | <0.001 |
| Bicho-Açu vs Curuá | | | | | 3.34 | 0.003 |
| Bicho-Açu vs Xamakorona | | | | | 6.09 | <0.001 |
| Curuá vs Xamakorona | | | | | 4.61 | 0.003 |

Abbreviations: df, degrees of freedom; SS, sum of squares; $R^2$, proportion of variance explained; *F*, PERMANOVA pseudo-*F* statistic.

**Supplementary Table 7. PERMANOVA of CAZyme beta diversity based on Bray–Curtis dissimilarities.** PERMANOVA results testing differences in CAZyme composition among Belgium (n = 39), Bicho-Açu (n = 31), Curuá (n = 20), and Xamakorona (n = 27). Pairwise post-hoc PERMANOVA contrasts are reported with *F* statistics and Benjamini–Hochberg FDR-adjusted *P* values. All tests used 999 permutations.

| **Comparison** | **Source** | **df** | **SS** | **$R^2$** | ***F*** | ***P*-adj** |
|---|---|---|---|---|---|---|
| Population | Model | 3 | 0.57 | 0.22 | 10.67 | <0.001 |
| | Residual | 113 | 2.01 | 0.78 | | |
| | Total | 116 | 2.58 | 1.00 | | |
| **PERMANOVA post-hoc test** | | | | | **F** | ***P*-adj** |
| Belgium vs Bicho-Açu | | | | | 16.35 | <0.001 |
| Belgium vs Curuá | | | | | 13.61 | <0.001 |
| Belgium vs Xamakorona | | | | | 18.47 | <0.001 |
| Bicho-Açu vs Curuá | | | | | 2.20 | 0.0226 |
| Bicho-Açu vs Xamakorona | | | | | 6.82 | <0.001 |
| Curuá vs Xamakorona | | | | | 5.43 | <0.001 |

Abbreviations: df, degrees of freedom; SS, sum of squares; $R^2$, proportion of variance explained; *F*, PERMANOVA pseudo-*F* statistic.
